# Heterochromatin protein 1α acts as an essential stabilizer of chromatin packing domains

**DOI:** 10.64898/2026.10.05.753292

**Authors:** Tiffany Kuo, Paola Carrillo Gonzalez, Nicolas Acosta, Lucas M. Carter, Emily Pujadas Liwag, Claudia V. Arriaga, Brittany Johnson, Cody L. Dunton, Luay M. Almassalha, Igal Szleifer, Vadim Backman

## Abstract

**Background:** The spatial organization of the human genome emerges from an enzyme-guided self-assembly process in which chromatin has a bidirectional relationship with transcription. Central to this architecture are chromatin packing domains, 50-200 nm structures exhibiting mass-scaling behavior. Nascent domains arise through transcription and cohesin/CTCF-mediated loop extrusion, while their maturation depends on conditions that permit differential access of epigenetic enzymes to their dense cores and continuously lower density periphery, resulting in the accumulation of heterochromatic marks in domain cores and euchromatic marks outside of the domain cores. In turn, packing domains support transcription by creating an “ideal zone” peripherally to their cores due to a nonmonotonic dependence of transcription rate on local chromatin density.

**Results:** We investigated the role of heterochromatin protein 1 alpha (HP1α), a key chromatin crosslinking factor with affinity to H3K9me3 marks, in maintaining packing domain organization. Paradoxically, short-term HP1α degradation resulted in overall downregulation of genes, contrasting with the canonical view that repression of heterochromatin allows chromatin to become more accessible for transcription. While some packing domains persisted, heterochromatic cores of dense packing domains were disrupted. When exposed to hypoxia, HP1α-depleted cells exhibited a dampened transcriptional response, showcasing that when domain maturation is disrupted, stress response pathways are not effectively activated.

**Conclusions:** This work highlights HP1α crosslinking as an essential component of domain maturation, as HP1α loss disrupts domain geometry and inhibits efficient transcriptional reprogramming. Our findings suggest an intriguing perspective on the potential relationship between chromatin architecture and gene regulation that could inform chromatin engineering-based treatments.

## Background

Higher-order genome organization is fundamentally characterized along two distinct yet complementary structural axes: genome geometry and genome topology. Genome geometry primarily focuses on the in-situ spatial arrangement of the chromatin polymer within a nucleus. Nanoscale imaging techniques, such as Chromatin Scanning Transmission Electron Microscopy (ChromSTEM) and Single-Molecule Localization Microscopy (SMLM), have demonstrated that the genome is organized into nanoscopic chromatin packing domains that promote gene regulation and overall genome stability^1,2^. These supranucleosomal structures are heterogeneous in size (50 to 200 nm) and density. Packing domains display power-law mass scaling behavior and have a continuous internal density gradient from a highly condensed core to a lower-density periphery^1,3,4^.

Genome topology mainly focuses on how topological features arise from the statistics of contact probabilities. Over the past two decades, understanding genome topology has been largely driven by proximity-ligation and sequencing-based connectivity methods, such as Hi-C^5,6^, ChIA-PET^7,8^, and GAM^9,10^. These techniques have revealed that statistically chromosomes partition into transcriptionally active “A” and inactive “B” subnuclear compartments. Within these genomic regions, there are topologically associated domains (TADs) that display high self-interaction frequencies^11^. The maintenance of three-dimensional chromatin organization has been strongly implicated in regulating essential cellular functions in the eukaryotic genome, including DNA damage repair, transcription, replication, cell division, and development^12,13^.

Rather than relying on a rigid hierarchical folding process, current paradigms suggest that these nanoscale packing domains emerge through a multimodal, enzyme-guided self-assembly process. Domain nucleation can be facilitated by both loop extrusion and active transcription, which generate forced chromatin returns that reconfigure the chromatin heteropolymer into volumes^4,14^. Following this initial nucleation, packing domains mature as a function of nucleosome compaction, molecular crowding, and the segregation of epigenetic enzymes biased by their molecular size and diffusivity^15,16^. Due to the mass fractal structure of packing domains, smaller (average size ∼2 nm) repressive epigenetic enzymes diffuse more readily into the denser domain cores than larger (average size ∼6 nm) euchromatic enzymes, resulting in the accumulation of repressive (H3K9me3/H3K27me3) marks in the domain core and euchromatic marks (H3K4me3/H3K27ac) in the domain periphery^17^. This power-law density gradient creates an “ideal zone” on the domain periphery that optimizes physicochemical conditions for transcriptional reactions^4^.

Packing domain geometry plays a critical role in regulating transcriptional processes^18–21^. More specifically, packing domain cores are fundamental structural components that support transcription of genes associated with packing domains. This highlights how transcriptional changes due to an external stimulus can be modulated by packing domains as well as the rate at which they form and mature.

Recent work has further revealed that introns and intergenic regions may generate volumes that position exons on the surfaces of packing domains^22^. Overall, these principles establish the strong potential of utilizing packing domains as physical geometric elements to regulate transcriptional memory and reprogramming.

Under this self-organizing framework, the structural integrity of the domain core becomes a critical prerequisite for transcription. Thus, we utilized RNA sequencing as well as nanoscale imaging modalities and analyses to study the role of heterochromatin protein 1 alpha (HP1α), a critical heterochromatic crosslinker, in the domain maturation process. We tested the hypothesis that degradation of a domain-stabilizing protein would result in the loss of H3K9me3 cores and disrupt the domain maturation process. We observed that loss of HP1α led to increased downregulation of transcriptional pathways. While some packing domains persisted, dense heterochromatic cores were readily disrupted. Moreover, loss of HP1α attenuates stress response pathways, suggesting that packing domain maturity plays a role in supporting optimal gene transcription. Our findings suggest a strong mechanistic link between chromatin architecture, gene regulation, and human diseases that can be utilized to enhance chromatin engineering-based therapies. Ultimately, this work establishes HP1α as a critical stabilizing protein that maintains the structural integrity of packing domain cores, enhancing our existing framework of the packing domain lifecycle.

## Results

### Short-term HP1α degradation leads to downregulation of transcriptional processes

We utilized human bone osteosarcoma U2OS cells that had the HP1α auxin-inducible degron for rapid and reversible depletion of the HP1α protein^23^. Using confocal microscopy and flow cytometry, we confirmed that HP1α fluorescence decreases after 5 hours of auxin treatment **(Fig. S1A-C)**. These results confirmed the efficient degradation of endogenous HP1α, enabling us to investigate its role in maintaining higher-order chromatin organization. We performed bulk RNA sequencing to test the effects of HP1α depletion on transcriptional regulation. Differential gene expression analysis revealed 15 significantly upregulated and 103 significantly downregulated genes in auxin-treated cells relative to control **(Fig. 1A)**. This decreased transcriptional response contrasts with the canonical expectation that degradation of a transcriptional repressor would lead to increased chromatin accessibility and subsequent transcription. This paradox led us to hypothesize whether loss of a crosslinking protein leads to compromised structural integrity of packing domains, specifically dense, heterochromatic cores that may serve as anchors of transcriptional memory.

**Figure 1:**
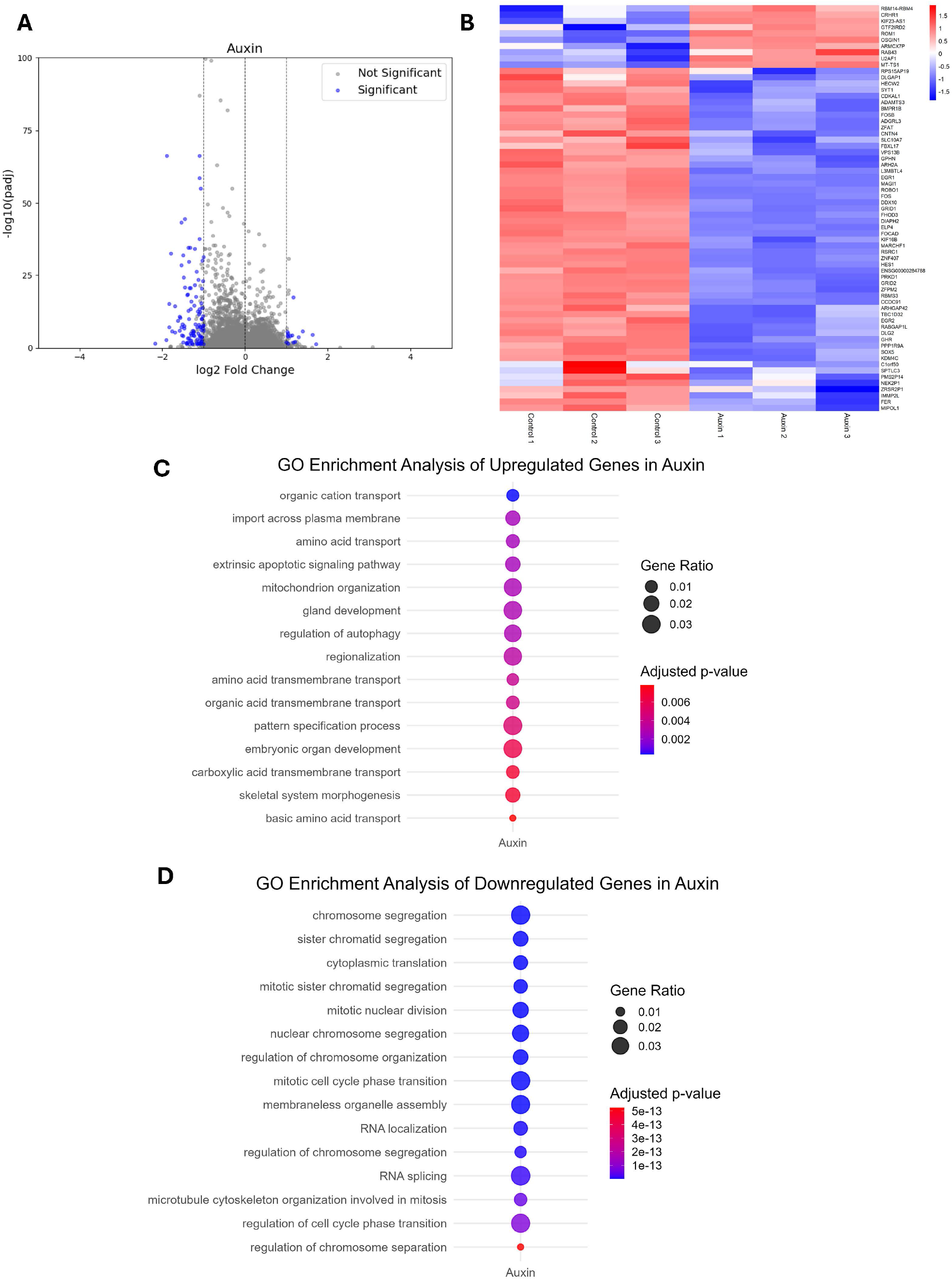
HP1α degradation leads to global downregulation of transcriptional processes. A. Volcano plot of differentially expressed genes (15 significantly upregulated and 103 significantly downregulated genes) after 5 hours of auxin treatment in U2OS-HP1α-AID cells. B. Heatmap of the top 65 differentially expressed genes in all three replicates of Control and Auxin conditions. C. Top 15 upregulated Gene Ontology (GO) biological processes (BP) terms in auxin treatment. D. Top 15 downregulated Gene Ontology (GO) biological processes (BP) terms in auxin treatment.

We identified the top 65 differentially expressed genes in auxin-treated osteosarcoma cells to determine whether downstream transcriptional programs were altered when HP1α is degraded. Immediate early response genes such as FOS/FOSB, EGR1/EGR2, and HES1 were differentially expressed, suggesting that HP1α depletion suppresses essential cell survival pathways. Auxin treatment also suppressed genes related to RNA processing and transcriptional regulation, such as U2AF1, ZNF407, and RSRC1 **(Fig. 1B)**. Furthermore, we conducted Gene Ontology (GO) enrichment analysis to uncover which biological processes were altered upon treatment. GO enrichment analysis revealed that HP1α depleted cells promoted processes related to organic cation/amino acid transport, mitochondrion organization, gland development, autophagy regulation, and regionalization **(Fig. 1C)**. Furthermore, HP1α depleted cells suppressed pathways related to chromosome segregation, mitotic cell cycle phase transition, membraneless organelle assembly, and RNA splicing **(Fig. 1D)**. Together, these results indicate that degradation of a heterochromatic crosslinker induces a transcriptionally repressive state in osteosarcoma cells, suggesting that HP1α loss induces alterations in higher-order chromatin organization.

For C-D, the sizes of the circles represent the proportion of differentially expressed genes that fall into a specific biological pathway, while the colors represent the statistical significance (adjusted p-value).

### HP1α maintains higher-order chromatin architecture by stabilizing heterochromatic cores of mature packing domains

We utilized Single Molecule Localization Microscopy (SMLM) for its multi-label capability to image epigenetic marks and transcriptional sites at sub-20 nm resolution to examine whether changes in the molecular composition of packing domains could explain the paradoxical transcriptional data. We labeled EdU to image DNA density and identify packing domains as well as H3K9me3 to serve as a proxy for packing domain cores^2,4^. This provided a metric for identifying more stabilized packing domains **(Fig. 2A)**. Using this two-color SMLM data of EdU-labeled DNA and H3K9me3, we investigated the association of H3K9me3 with EdU-defined packing domains. When HP1α was depleted, there was an increased percentage of packing domains that were not associated with heterochromatic cores (39.2% in control and 60.8% in auxin) **(Fig. 2B)**. This data demonstrates the weakened association between packing domains and dense, heterochromatic cores due to degradation of a crucial heterochromatic crosslinker.

**Figure 2:**
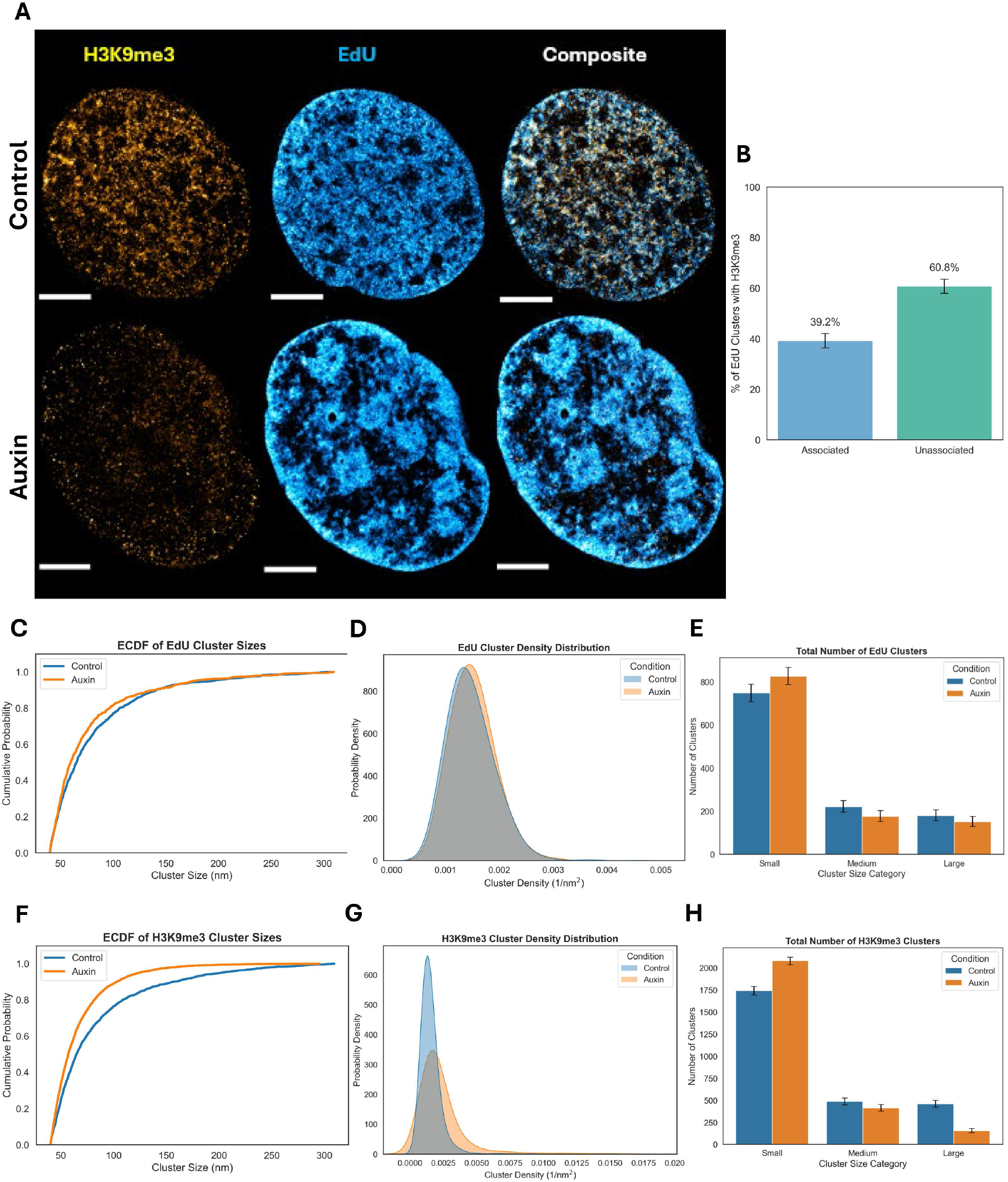
HP1α maintains higher-order chromatin architecture by stabilizing heterochromatic cores of mature packing domains. A. SMLM images of U2OS-HP1α-AID cells with and without auxin treatment, where H3K9me3 is in yellow and EdU is in blue. Scale bars = 5 μm. B. Percentage of EdU cluster sizes that had H3K9me3 marks associated with them in auxin-treated cells. 39.2% of EdU-defined packing domains were associated with H3K9me3 marks, while 68.2% of EdU-defined packing domains were not associated with H3K9me3. C. Empirical cumulative distribution function of EdU cluster sizes in control and auxin-treated U2OS-HP1α-AID cells, with auxin-treated cells having an increased number of smaller EdU-defined clusters. D. Probability density function of EdU cluster densities in control and auxin-treated U2OS-HP1α-AID cells shows that the density distribution of auxin treatment is shifted slightly to the right relative to the control. This suggests an increased tendency towards larger cluster densities in the auxin-treated cells. E. Total number of EdU-defined domains in each category size, where small domains are less than 80 nm, medium domains are 80-120 nm, and large domains are greater than 80 nm, in control and auxin-treated U2OS-HP1α-AID cells. F. Empirical cumulative distribution function of H3K9me3 cluster sizes in control and auxin-treated U2OS-HP1α-AID cells, with auxin-treated cells having an increased number of smaller EdU-defined clusters. G. Probability density function of H3K9me3 cluster densities in control and auxin-treated U2OS-HP1α-AID cells shows that the density distribution of auxin treatment has a higher and narrower peak relative to the control. This indicates a greater concentration of H3K9me3 cluster densities around 0.00125 and less variability relative to the control. H. Total number of H3K9me3-defined domains in each category size (small domains are less than 80 nm, medium domains are 80-120 nm, and large domains are greater than 80 nm) in control and auxin-treated U2OS-HP1α-AID cells.

To further uncover whether HP1α loss contributed to changes in the structural compositions of packing domains, we not only quantified EdU and H3K9me3 cluster sizes but also calculated localizations per cluster size, which serves as a proxy for cluster density. Cluster density provides additional insights into how molecular compositions of packing domains or heterochromatic cores change within a given area, revealing distinct structural properties that cannot be captured by cluster size alone. After auxin treatment, we observed a slight increase in the number of EdU-defined domains that were more uniform (approximately 50-60 nm in radii) and were slightly denser. We also saw an increased number of smaller EdU-defined packing domains (less than 80 nm) after auxin treatment **(Fig. 2C-D, S2A-C)**. Furthermore, the number of medium (80-120 nm) and large (greater than 120 nm) packing domains slightly decreased **(Fig. 2E)**. After auxin treatment, H3K9me3-labeled clusters had increased cluster sizes and decreased cluster densities **(Fig. 2F-G, S2D-F)**. When the H3K9me3 clusters were binned based on their sizes, we observed an increase in the number of smaller clusters, a decrease in the number of medium clusters, and a substantial decrease in the number of large clusters **(Fig. 2H)**. One potential explanation as to why there is a shift in the distribution of packing domain sizes is that larger structures are breaking down into smaller ones due to the loss of a crucial heterochromatic crosslinker. The substantial reduction in larger, heterochromatic clusters aligns with our hypothesis that HP1α acts as an essential stabilizer of dense packing domain cores. Together, these observations suggest that packing domains are unable to stabilize and fully mature without HP1α acting as a crucial crosslinking protein.

We plotted the distribution of the number of localizations within cluster sizes to demonstrate how molecular composition scales with cluster size, allowing us to better characterize the structural organization of nanoscale packing domains. In auxin-treated cells, the number of localizations in EdU clusters larger than 120 nm increased, but there were fewer points within the 200-300 nm range **(Fig. 3A)**. This indicates that there is a loss of larger packing domains, with a shift towards medium-sized packing domains, aligning with previous data showing a breakdown of larger structures when HP1α is degraded. In auxin-treated cells, the number of localizations in H3K9me3 clusters substantially increased in clusters less than 120 nm, with a subset exhibiting particularly high localization counts **(Fig. 3B)**. This increased number of localizations in smaller H3K9me3 clusters suggests that the structural integrity of packing domain cores (marked by H3K9me3) is heavily compromised due to HP1α depletion.

**Figure 3:**
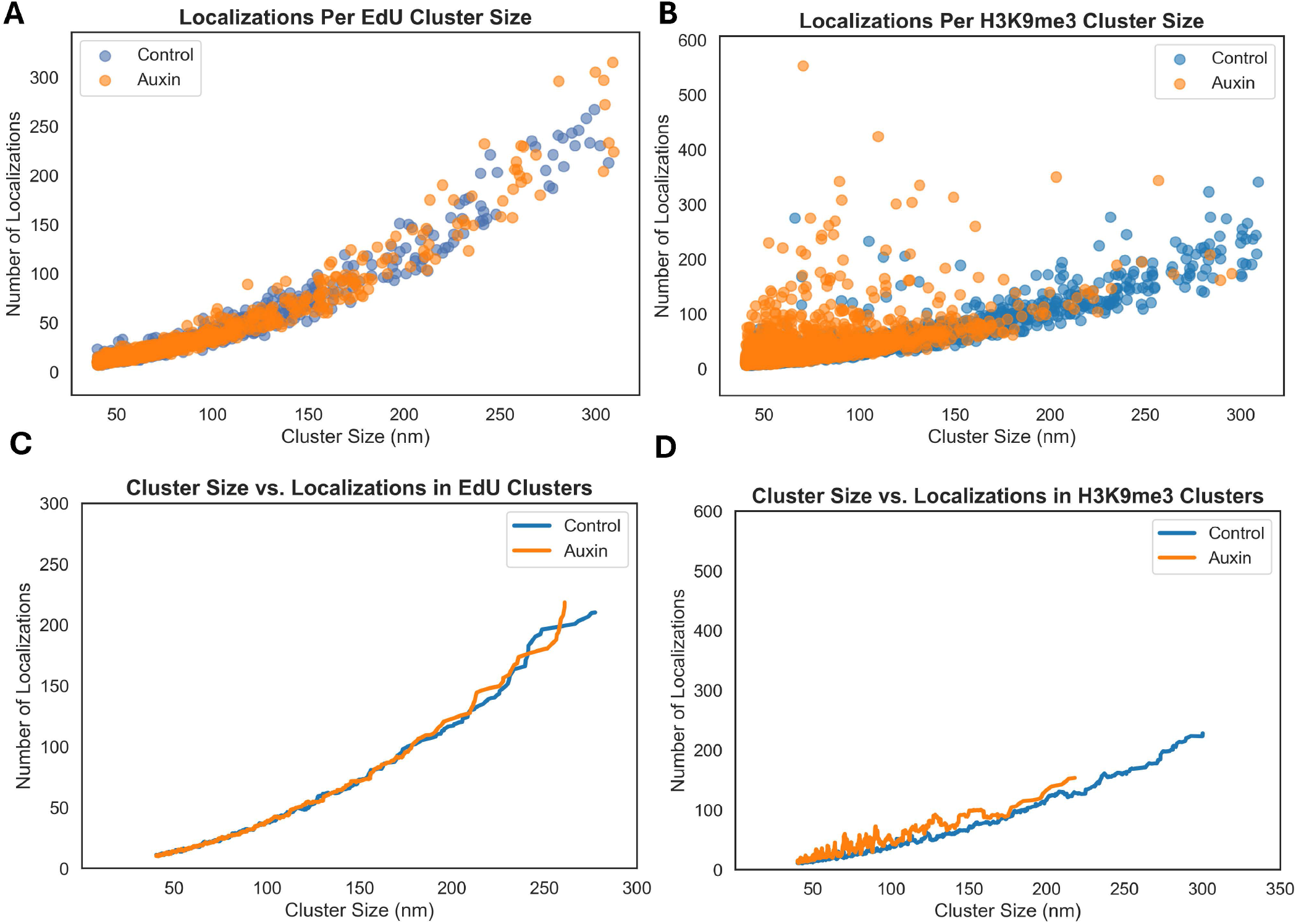
HP1α depletion disrupts spatial organization of mature packing domain cores. A. Quantification of the number of localizations per EdU cluster size in control and auxin-treated U2OS-HP1α-AID cells. B. Quantification of the number of localizations per H3K9me3 cluster size in control and auxin-treated U2OS-HP1α-AID cells. C. Moving average window profiles of the number of localizations across all EdU cluster sizes in control and auxin-treated U2OS-HP1α-AID cells. Curves were calculated using a sliding window of 20 nm. Both curves showed similar trends until around 200 nm, where the auxin trend line ends earlier relative to the control. D. Moving average window profiles of the number of localizations across all H3K9me3 cluster sizes in control and auxin-treated U2OS-HP1α-AID cells. Curves were calculated using a sliding window of 20 nm. The auxin curve had more variability and ended earlier at around 220 nm relative to the control.

We also generated moving average window profiles to capture the average number of localizations for clusters within a 20 nm window. This smooths out short-term fluctuations and highlights trends that may be obscured by how variable individual data points can be. In the EdU data, we observed similar trends in control and auxin conditions until approximately 200 nm, where the trend lines start to deviate from each other and the auxin trend line ends earlier relative to the control **(Fig. 3C)**. In the H3K9me3 data, the auxin curve had more variability and ended early at around 220 nm relative to the control **(Fig. 3D)**. Together, these findings suggest that HP1α loss prevents packing domains from fully maturing as there are fewer, larger packing domains. The increased abundance of small H3K9me3 clusters may reflect a breakdown of H3K9me3 cores as new packing domains are trying to form but cannot maintain this stability without a crucial crosslinking protein. The subset of clusters that have a greater number of H3K9me3 localizations may represent structures that have already acquired the essential stabilization mediated by histone modifications^24,25^, ionic forces^26–28^, and crosslinking proteins^29,30^. Since optimal gene transcription heavily relies on how stable packing domain cores are, loss of a critical crosslinking protein leads to compromised structural integrity of dense, heterochromatic cores, which is a key characteristic of mature packing domains^2,4^. As a result, the optimal regions that are essential for size-based nucleosome-modifying enzymes to access the core are diminished and transcriptional output is reduced, providing a potential explanation for the paradoxical transcriptional data.

### HP1α degradation destabilizes overall mesoscale chromatin organization

To substantiate the Single-Molecule Localization Microscopy data, we used Partial Wave Spectroscopic (PWS) microscopy to measure the statistical properties of chromatin structure due to its sensitivity to 20 nm chain sizes in live cells. With this technique, variations in nanoscopic-macromolecular structure are measured by analyzing the spectral dependence in light scattering from the packing of chromatin into higher-order structures, such as chromatin packing domains, while temporal variations in scattering allow measurement of chromatin mobility^31–33^. Since chromatin is organized into mass-fractal packing domains, average nuclear chromatin packing scaling (*D_n_*_)_ represents the fractal dimension of chromatin. This scaling factor is a statistical property of chromatin that determines the relationship between a chromatin chain and how it fits into three-dimensional space. Since chromatin is a complex polymer, the mass of chromatin contained within the domain *M* is related to the size of the domain *r* through the scaling exponent *D_n_*^1,3^. Within a packing domain, *D_n_* can theoretically range from 5/3 if the chromatin can exhibit a self-avoiding walk state, to 3 when chromatin is organized into a fractal globule state. However, polymers in a theta solvent, like chromatin, exhibit properties of a random walk model, which limits the range of *D_n_* to 2 < *D* < 3^5, 34,35^.

When HP1α is degraded upon the addition of auxin, there was minimal change in chromatin packing scaling in comparison to the control from 2.624 to 2.612, with a 95% CI [2.617, 2.631] and 95% CI [2.605, 2.619], respectively (p-value = 0.0197) **(Fig. 4A, B)**. Utilizing *D_n_* as a metric to assess overall chromatin organization between 20 and 200 nm, a marginal change in chromatin packing scaling suggests that the overall chromatin architecture largely remains intact despite HP1α loss. To further understand why there was a minimal effect, we utilized Dual-PWS to obtain dynamic measurements, as this technique captures variations in temporal interference from macromolecular movement. This allows for quantification of fractional moving mass (fraction of chromatin demonstrating coherent motion within a diffraction-limited volume)and effective diffusion coefficient (rate of motion at which chromatin segments move within the nucleus) without exogenous labels^3,33,36,37^. Despite the minimal change in higher-order chromatin structure that was observed with *D_n_*, there were much larger changes in fractional moving mass upon auxin treatment from 2.189 x 10^-19^ g to 2.586 x 10^-19^ g (18.1% increase) with a p-value < 0.0001 **(Fig. 4C)**. Effective diffusion coefficient substantially decreased from 0.004773 to 0.004362 μm^2^s^-1^ (8.6% decrease) with a p-value < 0.0001 **(Fig. 4D)**. With nuclear volume remaining the same, an increase in fractional moving mass correlates with the remaining structures being larger, or more mature. Moreover, a decrease in effective diffusion coefficient is consistent with how these larger packing domain structures are more mobile, but are restricted in a dense, nuclear environment. When paired with the super-resolution imaging data, this suggests that packing domains are constantly trying to form and mature but are unable to do so without HP1α. However, packing domains that already have pre-existing crosslinking mechanisms (including HP1α) remain in the system. Taken together, these complementary dynamics measurements reveal how short-term HP1α degradation destabilizes mesoscale chromatin organization, further demonstrating the role of HP1α in stabilizing packing domains to reach full maturity.

**Figure 4:**
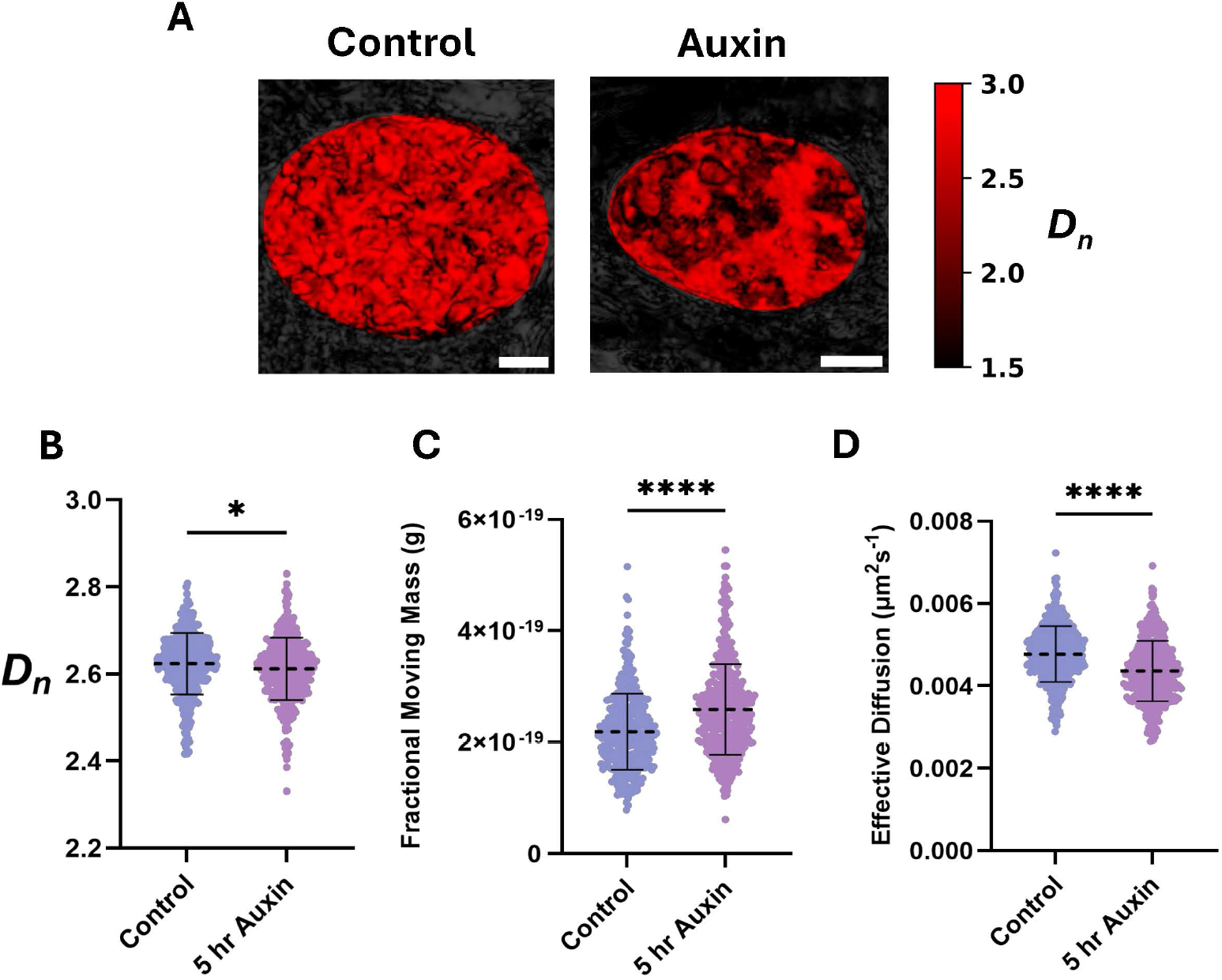
HP1α degradation destabilizes overall mesoscale chromatin organization. A. Representative images of control U2OS-HP1α-AID cells and U2OS-HP1α-AID cells treated with 5 hours of auxin. Brighter red indicates higher Dn (scale bars = 5 μm). B. Violin plots for chromatin packing scaling (Dn) in control U2OS-HP1α-AID cells and U2OS-HP1α-AID cells treated with 5 hours of auxin. C. Violin plots for fractional moving mass in control U2OS-HP1α-AID cells and U2OS-HP1α-AID cells treated with 5 hours of auxin. D. Violin plots for diffusion coefficient in control U2OS-HP1α-AID cells and U2OS-HP1α-AID cells treated with 5 hours of auxin.

For B-D: Data were compiled from three independent biological replicates. Significance was calculated by an unpaired t-test with Welch’s correction applied, with *P<0.05 and ****P<0.0001. (Control n=392; Auxin n=411)

### Mature packing domains retain higher structural integrity due to pre-existing crosslinking mechanisms

To further elucidate how crosslinking mechanisms play a role in packing domain stability and function, we applied our previously developed phenomenological model of domain formation to examine the differential sensitivities of nascent and mature packing domains in the absence of active transcriptional processes^4^. To distinguish between nascent and mature packing domains, we initialized our phenomenological model with parameter regimes corresponding to their experimentally observed structural properties. The initial chromatin packing state was described by the mass-radius scaling relation (M ∼ r*^D^*), where *D* specifies the initial packing state for nascent and mature packing domains^1,3^. Nascent and mature packing domains were distinguished by the relative contribution of biologically driven long-range returns. Active returns generated by transcription, RNA polymerase II loading, enhancer-promoter interactions, and cohesin-mediated loop extrusion nucleate local chromatin density pockets that give rise to packing domains. In contrast, entropic returns represent stochastic polymer contacts arising from confinement of the chromatin polymer and contribute to the background organization of chromatin but are insufficient to generate packing domains on their own **(Fig. S3A)**. Accordingly, nascent packing domains were initialized with a smaller fraction of active returns (10%) and a predominance of entropic returns (90%), whereas mature packing domains were initialized with a higher fraction of active returns (70%) and fewer entropic returns (30%) **(Fig. 5A)**. Following initialization, both domain types evolved under identical governing equations describing chromatin accumulation and packing reorganization **(see Materials and Methods).**

**Figure 5:**
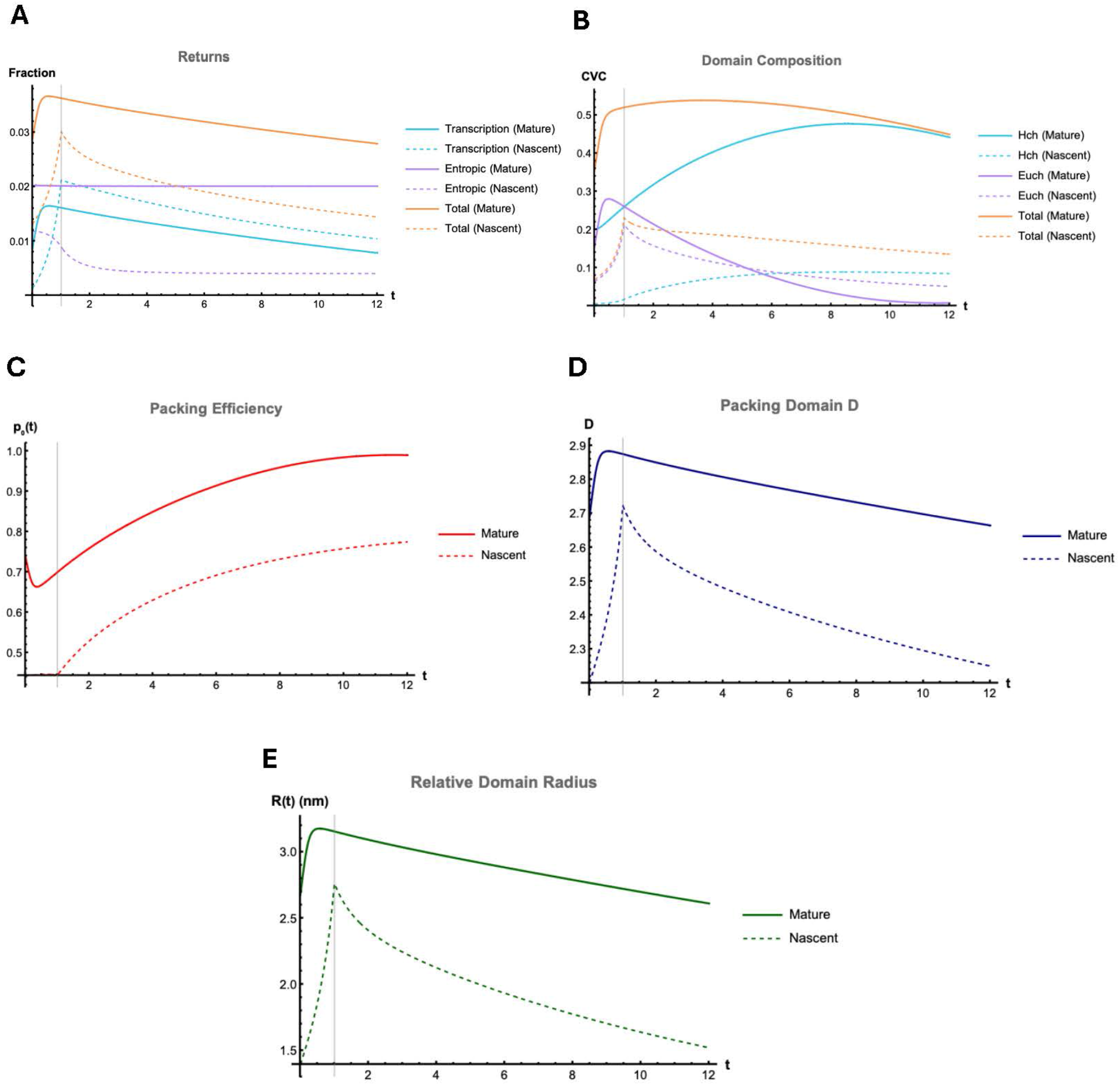
Model predictions show how mature domains retain higher structural integrity due to pre-existing crosslinking mechanisms. A. Phenomenological loop domain formation model predictions show that mature packing domains consist of fewer transcription-mediated returns and more entropic returns compared to nascent packing domains. B-E. Model predictions show mature packing domains retain high chromatin concentrations (CVC) (B), higher packing efficiency (C), higher packing domain *D* (D), and higher domain radius (E) compared to nascent packing domains when transcription was turned off after an hour.

When transcription was turned off after one hour, mature packing domains retained high chromatin volume concentration (CVC), indicating preservation of dense cores, whereas nascent packing domains failed to stabilize and did not accumulate sufficient heterochromatin to maintain domain integrity **(Fig. 5B)**. Mature packing domains also preserved higher packing efficiency, which measures how efficiently the chromatin fiber fills the packing domain volume, compared to nascent packing domains^1,4^. Similarly, mass-radius scaling *D_n_* and relative domain radius were retained in mature packing domains **(Fig. 5C-E)**. Together, these results demonstrate that mature packing domains maintain structural integrity even when transcription, an important boundary-forming process, is no longer active. This differential stability can be attributed to the presence of pre-committed structural returns in mature packing domains, which reinforce chromatin organization independently of ongoing transcription. These returns may arise from a combination of transcriptional history and non-transcriptional crosslinking mechanisms, including HP1α-mediated interactions^23^, ionic stabilization^28,38^, and macromolecular crowding^15^. In contrast, nascent packing domains lack sufficient pre-established returns and therefore fail to maintain stable packing once transcription is halted. Together, these results further enhance our perspective of a packing domain’s life cycle, highlighting how critical HP1α is in crosslinking domain cores for domain maturation, rather than creation, for mechanical long-term stability.

### Stress-induced transcriptional reprogramming was attenuated upon loss of HP1α crosslinking

Since we observed that packing domain cores were not stabilized without HP1α crosslinking, we wanted to test whether this inefficient domain maturation process would alter the cells’ ability to induce a beneficial transcriptional state in response to a stressor, such as cytotoxic stress, hypoxia, or heat shock. We hypothesized that HP1α-depleted cells would be unable to successfully induce a stress response due to the impaired domain maturation processes. To test this prediction, we pretreated cells with auxin for 5 hours (to destabilize domain cores and halt domain maturation processes) before exposing the cells to 24 hours of low-oxygen conditions. For simplicity, we will be referring to this treatment as “combination treatment.”

We performed additional bulk RNA sequencing experiments to assess whether halted domain maturation processes are associated with functional changes in transcriptional output. In hypoxia alone, we observed 333 significantly upregulated genes and 302 significantly downregulated genes **(Fig. 6A)**. Interestingly, in the combination treatment, we still observed transcriptional activity as 333 genes were significantly upregulated genes and 342 genes were significantly downregulated **(Fig. 6B)**. While both hypoxia and combination treatments resulted in the same number of significantly upregulated genes, the combination treatment led to more gene downregulation **(Fig. S4A)**. This aligns with our previous data demonstrating that loss of HP1α crosslinking prevents packing domains from maturing, resulting in sub-optimal conditions for transcription to occur. We also conducted Gene Ontology analysis to identify the pathways that were associated with each treatment. For hypoxia and combination treatments, there were similar upregulated biological processes, including regulation of autophagy, response to unfolded/misfolded proteins, RNA splicing, and tRNA/rRNA metabolic processes. Specifically, the enrichment of macromolecule methylation (slightly more significant in the combination treatment) suggests that cells are trying to survive by activating alternative methylation pathways **(Fig. S4B)**. Pathways related to chromosome segregation, nuclear chromosome segregation, mitotic cell cycle phase transition, membraneless organelle assembly, and nuclear division were downregulated in hypoxia and combination treatments **(Fig. S4C).**

**Figure 6:**
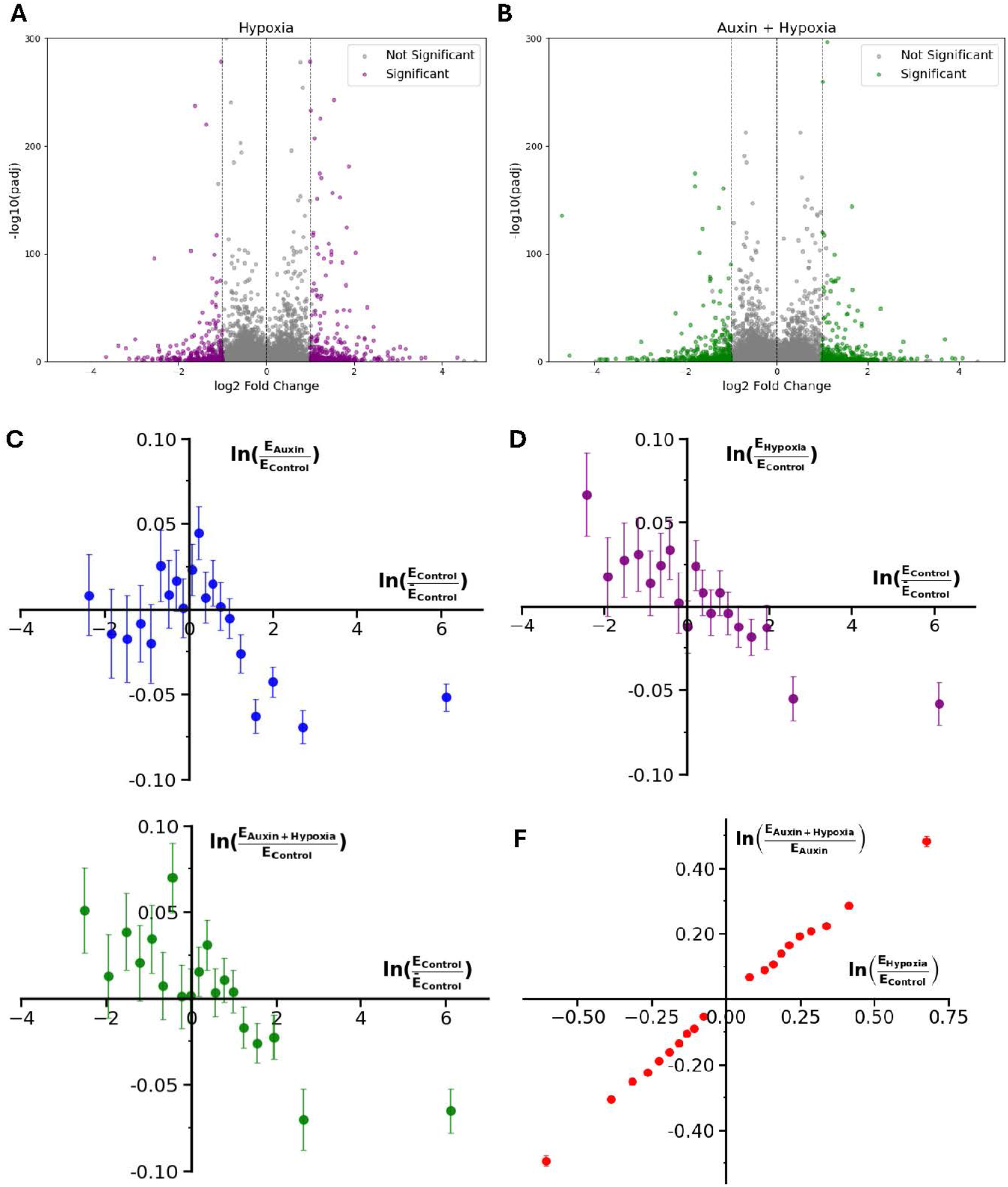
Stress-induced transcriptional reprogramming was attenuated upon loss of HP1α crosslinking. A. Volcano plot of differentially expressed genes (333 significantly upregulated and 302 significantly downregulated genes) after 24 hours of hypoxia treatment in U2OS-HP1α-AID cells. B. Volcano plot of differentially expressed genes (333 significantly upregulated and 342 significantly downregulated genes) after U2OS-HP1α-AID cells were pretreated with auxin for 5 hours before being exposed to 24 hours of hypoxia. C. In auxin-treated cells, a few initially downregulated genes are slightly downregulated, while a few unchanged genes are slightly upregulated. The majority of initially upregulated genes are further downregulated. D. In hypoxia-treated cells, initially downregulated genes are upregulated and initially upregulated genes are downregulated. E. In cells that were pre-treated with auxin before hypoxia, initially downregulated genes are upregulated and initially upregulated genes are downregulated with greater variability. F. Genes that were upregulated by hypoxia treatment remained upregulated in the combination treatment, and genes that were downregulated by hypoxia are still generally downregulated.

In order to evaluate the data quality and sample reproducibility, we also conducted principal component analysis (PCA) and Pearson correlation analyses on the bulk RNA sequencing data. PCA demonstrated that the biological replicates clustered closely together while treatment groups were clearly separated along both principal components **(Fig. S5A)**. Sample Pearson correlation analysis complemented this analysis by showing that there were high within-group correlations and hierarchical clustering of samples based on treatments **(Fig. S5B)**. This data confirmed that the three biological replicates were consistent and that the treatment effects contributed to the variation in the dataset.

To further elucidate the functional consequences when HP1α is depleted, we hypothesized that packing domain structures had a role in generating the phenotypes we observed. Dual-PWS microscopy revealed that cells exposed to hypoxia induced a modest increase in *D_n_*, rising from 2.592 to 2.646, with a 95% CI [2.583, 2.601] and 95% CI [2.637, 2.655], respectively (p-value < 0.0001). This suggests that nascent structures are being formed in response to a transcriptional reprogrammer. When HP1α was degraded for 5 hours before 24 hours of hypoxia, *D_n_* increased (albeit to a lesser extent than hypoxia) compared to the control, from 2.592 to 2.615, with a 95% CI [2.583, 2.601] and 95% CI [2.608, 2.622], respectively (p-value=0.0001) **(Fig. S6A, B)**. This indicates that even though mature packing domains cannot be formed, packing domains remain in the system. Consistent with previous data, these structures are most likely packing domains that are at the early stage of their life cycle and trying to stabilize. Both hypoxia and combination treatments resulted in increased fractional moving mass and decreased effective diffusion. The observed increase in the amount of chromatin present in both treatments is consistent with cells having a greater fraction of chromatin contained in new domains. Moreover, the decrease in effective diffusion coefficient correlates with these packing domain structures being mobile but restricted by the crowded nuclear environment **(Fig. S6C, D)**. Together, these complementary dual-PWS measurements suggest that, even though packing domains cannot reach full maturity, the system can still generate nascent structures to adapt to an external stressor.

To better characterize this dynamic relationship between domain maturation state and transcriptional outcomes, we also conducted more in-depth analyses of this RNA sequencing data. We grouped differentially expressed genes (p-value < 0.05) into quantiles based on their initial expression in control cells. This methodology allowed us to identify genes that had significant changes in expression upon treatment. We can interpret these plots in the context of packing domains by viewing the x-axis as a proxy for domain maturity, with the left side corresponding to nascent or smaller domains, the middle corresponding to intermediate domains, and the right side corresponding to more mature domains. These are based on our understanding of how packing domains are related to transcription – genes that are not being transcribed or are lowly transcribed are most likely not in domains or newly forming domains, while transcribed genes are in domains.

In auxin treatment, a few initially downregulated genes were slightly downregulated while a few unchanged genes were slightly upregulated. However, the majority of initially upregulated genes were further downregulated **(Fig. 6C)**. This suggests that immature and intermediate domains are not as affected by auxin treatment since these have not fully acquired heterochromatic marks, which are essential for HP1α crosslinking. On the other hand, mature domains are more affected by auxin treatment, as loss of a critical crosslinker induces loss of the ideal zones and subsequently decreases transcriptional efficiency. In hypoxia treatment alone, initially downregulated genes were upregulated and initially upregulated genes were downregulated **(Fig. 6D)**. Here, nascent domains are upregulated to transcribe hypoxia-specific stress response genes. Mature domains are downregulated as previously essential genes (before the hypoxia stressor is induced) are no longer needed. When we looked at cells that were initially destabilized via auxin treatment and then exposed to hypoxia, initially downregulated genes were upregulated and initially upregulated genes were downregulated with greater variability **(Fig. 6E)**. Despite loss of HP1α crosslinking, nascent packing domains can still acquire other enzymes, divalent ions, and crosslinking proteins to increase their domain-forming capacity. Due to the variability, we delved deeper into the data by looking at genes that were differentially expressed in hypoxia relative to control (x-axis) graphed against genes that were differentially expressed in the combination treatment relative to auxin (y-axis). Here, we see that genes upregulated by hypoxia were still generally upregulated in the combination treatment, and genes downregulated by hypoxia were still generally downregulated **(Fig. 6F)**. Since packing domains were demonstrated not to have been completely lost in the dual-PWS data, this observation suggests that the stressor can drive the re-formation and stabilization of existing packing domains, resulting in a similar transcriptional response. Interestingly, the slope of this trend is approximately 0.7599, suggesting that degradation of HP1α largely preserves transcriptional activation in response to low oxygen conditions but lowers its magnitude by 24%.

To further probe whether differentially expressed genes were associated with weak or strong packing domains before transcriptional reprogramming, we identified hypoxia-responsive genes that were significantly altered by hypoxia treatment relative to the control. We then binned these genes into three groups based on their initial expressions and plotted the average response of each group **(Fig. S7A)**. Genes that had low initial expression were upregulated by hypoxia when HP1α was present and remained upregulated by hypoxia after HP1α degradation. Genes that had medium initial expression were slightly upregulated by hypoxia but exhibited a slightly decreased response when exposed to hypoxia after HP1α degradation. Finally, genes that had high initial expression were downregulated by hypoxia with and without HP1α degradation. Overall, these results suggest that induction of the hypoxia response is largely preserved in genes that were initially lowly and highly expressed. Interestingly, genes in the medium initial expression group exhibit minimal change after auxin treatment. Since this graph plots the average response of each group, we also calculated the slopes of all genes within each binned group **(Fig. S7B)**. Here, we observed that the slopes of all binned groups were less than 1, suggesting that the hypoxia response is reduced after initial packing domain destabilization via HP1α degradation, aligning with previous observations of attenuated transcriptional response. We see that genes that were initially highly expressed retain the hypoxia response slightly better than the low-or medium-expression genes. These results point to a potential hypothesis that initial gene expression may influence domain maturation state.

For C-E: Change in gene expression of the treatment was plotted as a function of the initial gene expression level. Differentially expressed genes were then grouped into quantiles based on their expression levels in control cells. The average gene expression for control is plotted on the x-axis, while the change in gene expression after the corresponding treatment is plotted on the y-axis. Error bars are standard errors of the mean.

### Impaired domain maturation processes due to HP1α loss induce inefficient transcriptional pathways

To further understand the functional consequences of inefficient domain maturation processes, we measured nascent RNA production via 5-ethynyl uridine (EU) staining for 1 hour. In control cells, nascent RNA localizations were evenly distributed throughout the nucleus. In auxin-treated cells, nascent RNA localizations were more concentrated, resulting in the appearance of brighter puncta **(Fig. 7A)**. When the number of nascent RNA localizations was quantified per nuclear area, we saw that there were around 133 EU localizations per nuclear area in control cells and 163 EU localizations per nuclear area in auxin-treated cells **(Fig. 7B)**. These results suggest that disruption of packing domain geometry can result in activation of transcriptional activity due to increased accessibility throughout the nucleus.

**Figure 7:**
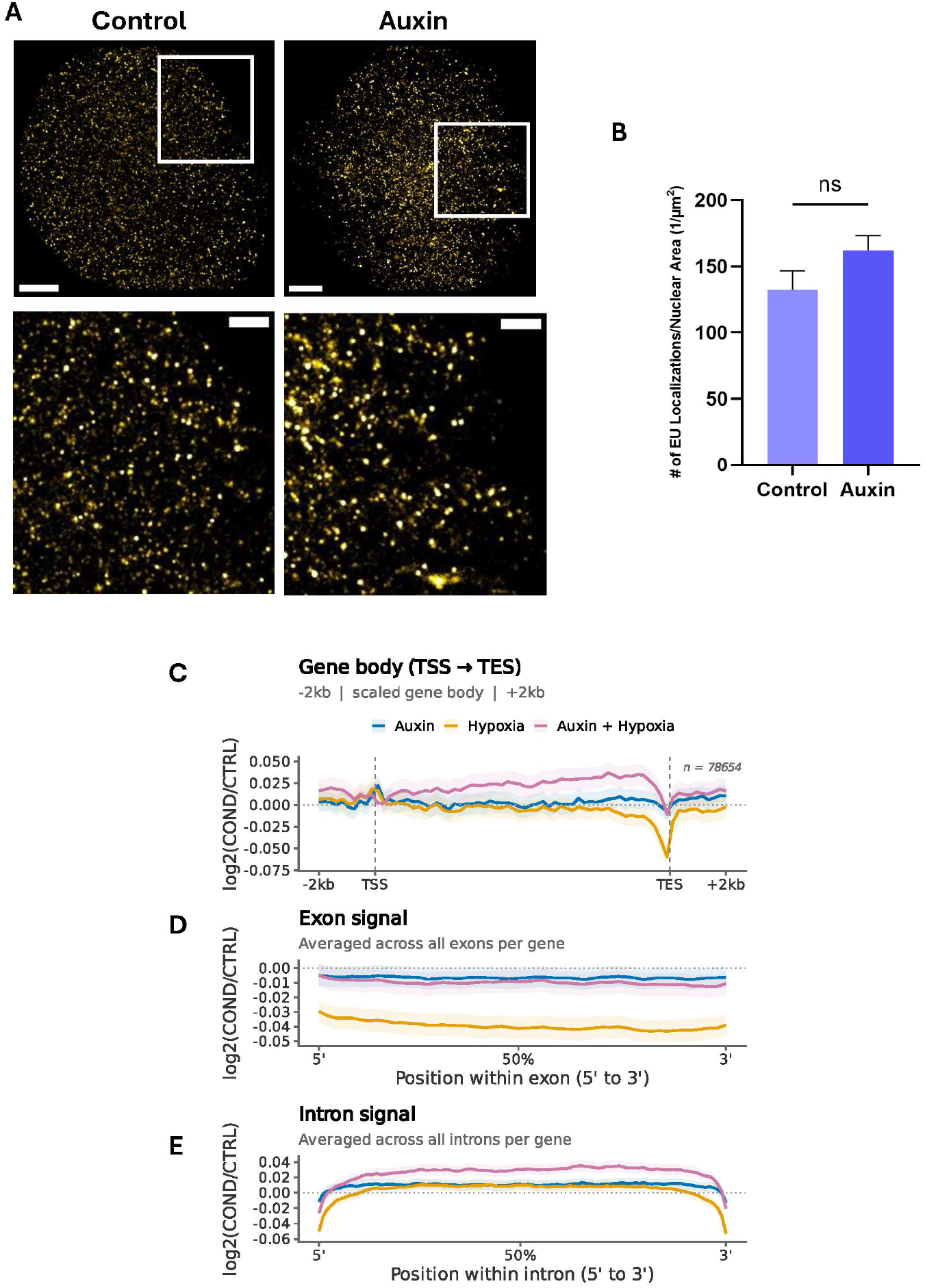
Impaired domain maturation processes due to HP1α loss induce inefficient transcriptional pathways. A. SMLM of nascent RNA measured by EU synthesis (yellow) in control and auxin-treated U2OS-HP1α-AID cells. Scale bar = 3 µm, Scale bar (zoom) = 1 µm. B. Quantification of the number of EU localizations per nuclear area in control and auxin-treated U2OS-HP1α-AID cells. Significance was calculated by an unpaired t-test with Welch’s correction applied. C. Plot of transcriptional signal from Transcriptional Start Site (TSS) to Transcriptional End Site (TES) in auxin, hypoxia, and combination treatments relative to the control. Region ± 2 kilobases. D. Plot of aggregate exon signal (ideal zone) in auxin, hypoxia, and combination treatments relative to the control. E. Plot of aggregate average intron signal (total volume) in auxin, hypoxia, and combination treatments relative to the control.

With prior work demonstrating that introns may generate volumes that position exons on the surfaces of packing domains, we also utilized the exon-to-intron (E/I) ratio as a proxy for the surface area/volume ratio of packing domains^22^. Hence, we calculated exonic and intronic contents across all conditions analyzed in the RNA sequencing dataset. When we quantified the average transcriptional signal across all genes after aligning them from the Transcription Start Site (TSS) to the Transcription End Site (TES), the profiles were comparable between auxin and hypoxia treatments, with a slight reduction in coverage near the TES in the hypoxia treatment. In contrast, the combination treatment coverage from the TSS to the TES was more abundant relative to auxin and hypoxia treatments alone, which is consistent with enhanced transcriptional signal **(Fig. 7C)**. This reduced transcriptional signal at the 3’ end may be due to exposure to a stressor that causes RNA polymerase to become less stable^39^. The enhanced transcriptional signal associated with the combination treatment may reflect the cells activating transcriptional reprogramming in response to a stressor, aligning with previous data suggesting that existing packing domains can re-form and stabilize despite loss of HP1α crosslinking. When we further examined exonic and intronic coverage, we observed that hypoxia treatment exhibited an increased exonic signal, which corresponds to a higher E/I ratio (higher surface area/volume ratio) **(Fig. 7D)**. This exposed surface area allows for increased accessibility of ideal zones for transcriptional reactions to occur, which is consistent with the system’s need to enhance hypoxia-specific transcriptional pathways. Furthermore, the combination treatment resulted in an increased intronic signal, meaning that packing domains exhibited a low E/I ratio (low surface area/volume ratio) when HP1α was depleted before hypoxia treatment **(Fig. 7E)**. This observation suggests that introns that are normally buried within packing domains are now exposed to transcriptional machinery. However, the system’s ability to transcribe functional products, such as hypoxia-responsive genes, is effectively suppressed due to packing domains being unable to properly mature. This further highlights how HP1α loss disrupts the essential structural elements of packing domains, specifically the cores, to provide the foundation for productive stress response pathways.

## Discussion

Uncovering the dynamic relationship between 3D genome organization and transcriptional regulation is particularly crucial to understanding the underlying mechanisms of processes such as DNA damage repair pathways, transcription, replication, cell division, and development^12,13,40^. In this study, we revealed how heterochromatin protein 1 alpha plays a crucial role in stabilizing packing domains and supporting effective transcriptional reprogramming. Loss of HP1α led to a paradoxical increase in downregulated genes, suggesting that packing domain cores were heavily disrupted **(Fig.1)**. Consistent with this prediction, we observed the overall distribution shift towards smaller packing domain sizes and a substantial reduction in large, heterochromatic clusters **(Fig. 2)**. While some large, dense structures remained, the increased abundance of small H3K9me3 clusters may reflect a breakdown of packing domain cores that cannot be properly stabilized **(Fig. 3)**.

Even though chromatin organization at the sub-200 nm length-scale largely remained similar after HP1α degradation, increased fractional moving mass and a lower effective diffusion coefficient suggested that larger structures remained in the system but were restricted in the dense, nuclear environment **(Fig. 4)**. This observation aligns with more stable packing domains persisting in the system, as nascent structures cannot transition into mature ones without HP1α. Simulations further reveal HP1α as an essential factor in stabilization, not creation, of packing domains, as mature packing domains that have pre-committed structural returns retain higher structural integrity **(Fig. 5)**.

To better characterize HP1α’s role in the domain maturation process, we used hypoxia as a model system of transcriptional reprogramming. We investigated how the genome responds to an external stimulus when packing domain maturation was disrupted and whether this created a new transcriptional state. When cells were pre-treated with auxin before hypoxia, the transcriptional response was similar to that of hypoxia alone. Since packing domain structures were not completely destroyed, this indicated that the transcriptional stimulus could drive existing packing domains to try to re-form and stabilize **(Fig. 6)**.

Nascent RNA synthesis increased with HP1α loss, suggesting increased accessibility throughout the nucleus due to impaired packing domain geometry. Increased intronic signal associated with the combination treatment further corroborates how HP1α loss disrupts the essential structural elements of packing domains, specifically the cores, which are crucial in initiating productive stress response pathways **(Fig. 7)**.

Overall, our findings demonstrate that nascent packing domains cannot mature as efficiently without HP1α providing essential stabilization of packing domain cores. In turn, the system has a lower transition rate into a new transcriptional state when exposed to an external stimulus. The observed paradoxical decrease in transcription upon HP1α degradation aligns with evidence showing that loss of heterochromatin-associated proteins or complexes can result in both gene upregulation and downregulation. Loss of PRC2, a chromatin-modifying complex that catalyzes H3K27me3, was associated with downregulation of genes that are specific for medium spiny neurons^41^. The timeframe at which heterochromatic marks are acquired is also important to note here. Prior work has demonstrated that upon 24 hours of GSK343 treatment, which inhibits EZH2 activity and reduces H3K27me3 activity, H3K9me3 intensity increased at the nuclear periphery relative to the control in HCT116 cells^42^. To complement this data, changes in nascent RNA production were also seen in HCT116 cells upon 24 hours of GSK343 treatment^4^. These findings showed that within 24 hours, there were substantial changes in heterochromatin (specifically H3K9me3) formation and degradation, demonstrating that this shorter timeframe can be utilized to investigate loss of a heterochromatin crosslinker on overall chromatin organization and dynamics. However, one of the limitations is the auxin degron system, as residual HP1α may persist after short-term protein degradation. Differences in cell cycle stages may also contribute to the observed heterogeneity in packing domain size and organization. Since nanoscale imaging techniques cannot capture dynamic processes, it remains an open question as to whether destruction of existing domains or impaired domain maturation is the primary driver of the observed phenotypes in this study.

Future studies that combine chromatin scanning transmission electron microscopy (ChromSTEM) with labeling for specific heterochromatic markers will further resolve how HP1α loss reshapes chromatin architecture at a higher resolution^1^. In parallel, gene-specific spatial mapping using resolution after single-strand exonuclease resection (RASER)-FISH or DNA-PAINT together with SMLM may also provide deeper insights into how hypoxia-specific genes are positioned within packing domains before and after perturbations^43–45^. By employing these gene-specific labeling techniques, we can further characterize how 3D packing domain architecture influences transcription. Additional time-course experiments can be conducted in HP1α-depleted cells where transcriptional responses will be measured at multiple time points after hypoxia treatment. This will provide additional insights into the rate of stress-induced transcriptional reprogramming when domain maturation processes are halted, further underscoring how packing domain geometry facilitates the transition between transcriptional states.

With this new perspective on HP1α’s role in chromatin organization and mechanics, several opportunities emerge for potential novel cancer therapeutics. Enhancing HP1α could impede chemotherapy resistance by slowing down or even preventing chemoresilient cancer types from transitioning into pro-survival transcriptional states^46,47^. Alternatively, dysregulation of packing domain geometry via HP1α degradation may limit the ability of cancer cells to undergo adaptive transcriptional programming that is commonly associated with drug resistance, improving the efficacy of chemotherapeutic drugs^48,49^. This highlights the strong potential of modulating HP1α activity to overcome chemoresistance. Ultimately, this study reveals that HP1α is not only a crucial stabilizing factor of packing domain geometry but also a regulator of adaptive transcriptional reprogramming.

## Conclusions

Our results demonstrated that HP1α acts as an essential stabilizer of packing domains, as new packing domains cannot mature efficiently. Within short-term HP1α depletion, we observed an increased number of downregulated genes, with differentially expressed genes primarily being immediate early response genes. While some large, dense structures remained, the increased abundance of small H3K9me3 clusters reflected a breakdown of packing domain cores that could not be stabilized. Simulations further characterized HP1α as a factor in packing domain stabilization, rather than creation. When the domain maturation process was disrupted, the hypoxia-induced stress response remained intact but was attenuated, suggesting that although cells retain the ability to respond to a stressor, efficient transcriptional reprogramming heavily relies on the formation and stabilization of packing domains.

Together, this work highlights how HP1α is not only a structural protein that binds to constitutive heterochromatin marks but also an essential crosslinker of packing domain cores that contributes to the overall stability of higher-order chromatin organization and function. Future work on uncovering the roles of other HP1 isoforms and regulatory proteins may reveal deeper insights into the complex interplay between domain geometry and transcription, providing additional targets in clinical settings.

## Materials and Methods

### Cell Culture

U2OS HP1α-AID cells were grown in McCoy’s 5A Modified Medium (#16600-082, Thermo Fisher Scientific, Waltham, MA) supplemented with 10% Fetal Bovine Serum (#16000-044, Thermo Fisher Scientific, Waltham, MA) and penicillin-streptomycin (100 μg/ml; #15140-122, Thermo Fisher Scientific, Waltham, MA). The U2OS HP1α-AID-sfGFP degron clone was created as previously described in Strom et al^23^. All cells were cultured under recommended conditions at 37°C and 5% CO_2_ and maintained between passages 5 and 20. Cells were allowed at least 24 hours to re-adhere and recover from trypsin-induced detachment prior to imaging or treatment. All imaging was performed when the surface confluence of the dish was between 40–70%.

### Treatments

For auxin treatment, U2OS-HP1α-AID-sfGFP cells were plated at 50,000 cells per well of a 6-well plate (Cellvis, P12-1.5H-N). Cells were given at least 24 hours to re-adhere prior to treatment. For all experiments, 500 mM Indole-3-acetic acid sodium salt (IAA, Sigma Aldrich, #I28860) was solubilized in UltraPure DNase/RNase-Free Distilled Water (#10977015). Optimal auxin treatment time was determined based on results from Strom et al.^23^ as well as additional immunofluorescence and flow cytometry experiments **(Fig. S1A-C)**. For hypoxia treatments alone, cells were exposed to low concentrations of oxygen at 0.5% for 24 hours before subsequent imaging and sequencing. For combination treatments of auxin and hypoxia, cells were exposed to 5 hours of IAA before being exposed to low concentrations of oxygen at 0.5% for 24 hours before subsequent imaging and sequencing.

### Fixed-Cell Flow Cytometry (FACS) Analysis

Flow cytometry analysis for U2OS-HP1α-AID-sfGFP cells to determine proper auxin treatment concentration was performed on a BD LSRFortessa Cell Analyzer FACSymphony S6 SORP system, located at the Robert H. Lurie Comprehensive Cancer Center Flow Cytometry Core Facility at Northwestern University in Evanston, IL. For all FACS analysis, the same protocol was used. After auxin treatment, cells were harvested and fixed. Briefly, cells were washed with DPBS (Gibco, #14190-144), trypsinized (Gibco, #25200-056), neutralized with media, and centrifuged at 500 x g for 5 minutes. Cells were then resuspended in 500 μL of 4% PFA and DPBS and fixed for 10 minutes at room temperature, followed by centrifugation and resuspension in cold FACS buffer (DPBS with 1% BSA and 2mM EDTA added) at 4°C until analysis could be performed. Flow cytometric analysis (FACS) data were analyzed using FlowJo software version 10.6.1.

### Immunofluorescence Sample Preparation

U2OS HP1α-AID-sfGFP cells were plated at 50,000 cells per well of a 6-well plate (Cellvis, P12-1.5H-N). Following auxin treatment, cells were washed twice with 1 × Phosphate Buffered Saline (PBS) (Gibco, #10010031). Cells were fixed with 4% paraformaldehyde (PFA) (Electron Microscopy Sciences, #15,710) for 10 min at room temperature, followed by washing with PBS 3 times for 5 min each. Cells were permeabilized using 0.2% TritonX-100 (10%) (Sigma-Aldrich, #93443) in 1 × PBS, followed by another wash with 1 × PBS 3 times for 5 min each. Cells were blocked using 3% BSA (Sigma-Aldrich, #A7906) in PBST (Tween-20 in 1 × PBS) (Sigma-Aldrich, #P9416) at room temperature. The following primary antibody was added overnight at 4°C: anti-HP1α (ab109028, dilution 1:250). Cells were washed with 1 × PBS 3 times for 5 min each. Cells were washed with 1 × PBS 3 times for 5 min each. The following secondary antibody was added for 1 hour at room temperature: Invitrogen Goat anti-Rabbit IgG (H + L) Highly Cross-Adsorbed Secondary Antibody, Alexa Fluor 647 (Thermo Fisher Scientific, #A-21245, dilution 1:200). Cells were washed with 1 × PBS 3 times for 5 min each. Finally, cells were stained with DAPI (Thermo Fisher Scientific, #62248, diluted to 0.5 μg/mL in 1 × PBS) for 10 min at room temperature. Prior to imaging, cells were washed with 1 × PBS twice for 5 min each.

### Confocal Microscopy Imaging

Confocal images were obtained using the Nikon SoRa Spinning Disk confocal microscope located at the Biological Imaging Facility at Northwestern University in Evanston, IL (RRID: SCR_017767). The Nikon Ti2 inverted microscope is equipped with a Yokogawa CSU-W1 dual-disk spinning disk with 50 μm pinholes, a 50 μm pinhole with micro-lenses SoRa disk, and an ORCA-Fusion Digital CMOS camera (Hamamatsu). Images were collected using a 60x (1.42 NA) oil-immersion objective lens. sfGFP was excited with a 488-nm laser, Alexa Fluor 647 was excited with a 638-nm laser, and DAPI was excited with a 405-nm laser. All imaging data were acquired by Nikon NIS Elements acquisition software.

### RNA-Seq Library Preparation

Total RNA extraction was performed using the RNeasy Mini Kit (Qiagen, #74106) following the manufacturer’s protocol. The conditions for these samples were control, 5-hour auxin, 24-hour hypoxia, and 5-hour auxin followed by 24-hour hypoxia. These samples were collected with three biological replicates per condition. RNA-Seq libraries were prepared using the Illumina TruSeq Stranded mRNA Library Prep Kit. Library quality was measured using the Qubit 2.0 and Bioanalyzer. Bulk mRNA sequencing was conducted at the Northwestern University NUSeq Core Facility. Each sample was sequenced to a depth of 20-25 million reads using NovaSeq X Plus (50 bp read length).

### RNA-Seq Data Analysis

RNA-seq reads were preprocessed with FastQC and aligned with STAR to the human reference genome GRCh38 (Ensembl). Aligned samples were converted to binary alignment map (BAM) files, which were then normalized and compared for differential gene expression using the DESeq2 package in R. Volcano plots were generated using the scatterplot function in Python, and the heatmap generated using the pheatmap package in R. To identify significantly upregulated genes, we set a threshold of log2 Fold Change greater than or equal to 1 and an adjusted p-value less than 0.05. Similarly, to identify significantly downregulated genes, we set a threshold of log2 Fold Change less than or equal to 1 and an adjusted p-value greater than 0.05. We used the clusterProfiler package in R to perform Gene Ontology enrichment analysis.

For the Exon/Intron analysis, raw FASTQ files were quality trimmed using TrimGalore (v0.6.10)^50^ with default single-end parameters and automatic adapter detection. Trimmed reads were aligned to the human reference genome (GRCh38) using HISAT2 (v2.1.0)^51^ with splice-aware alignment (--dta flag). Aligned reads were filtered using SAMtools (v1.14)^52^ to remove unmapped, secondary, supplementary, low-quality, and duplicate-flagged reads (flag-F 3852, MAPQ ≥ 30). Filtered reads were converted to BAM format, sorted, and indexed using SAMtools. Alignment rates and flagstat summaries were computed per sample for quality control.

Per-replicate filtered, sorted BAMs were merged per condition using *samtools merge*, then re-sorted and indexed. Merged BAMs represent all three replicates per condition combined. Library size scale factors were computed from the merged BAMs to normalize for differences in sequencing depth between conditions. The total number of mapped reads was counted per condition using *samtools view –c –F 4*.

Scale factors were calculated as Control mapped reads / condition mapped reads, anchoring all conditions to the Control library size (SF_Control = 1.0, SF_Auxin = 0.930660, SF_Hypoxia = 0.807493, SF_Auxin_Hypoxia = 0.938752).

BigWig files were generated per condition using deepTools (v3.5.1):^53^ Log2 ratio bigWigs — pairwise log2(condition/Control) tracks generated using *bamCompare* with –-operation log2, –-binSize 20, –-smoothLength 60, –-ignoreDuplicates, and –-pseudocount 1. Scale factors for both the condition and Control BAMs were applied simultaneously via –-scaleFactors to ensure depth-normalized comparison. These tracks were used for all downstream metagene analyses.

Metagene profiles were computed in R using rtracklayer^54^ and plyranges^55^ to extract log2 ratio bigWig signal over three interval types: gene bodies (TSS to TES with ±2kb flanks), exon intervals, and intron intervals. Gene annotations were derived from the Ensembl GRCh38 release 115 GTF using GenomicFeatures::makeTxDbFromGFF()^56^.

Gene body profiles used a scale-regions approach with 50 bins across the gene body, 10 bins over the ±2kb upstream and downstream flanks, respectively (70 bins total). Exon profiles used 50 bins per individual exon interval; intron profiles used 100 bins per intron interval. Intervals shorter than the bin count were excluded. Signal was extracted per bin using weighted overlap averaging against the RleList representation of each bigWig. Mean signal and standard error were computed across all genes per condition per bin position.

Three plots were generated: (1) all expressed genes (TPM ≥ 1) combined, (2) stratified by expression tertile (Low/Unexpressed, Medium, High) computed from RNA-seq TPM values, and (3) stratified by genomic exon/intron ratio (E/I) tertile (Low E/I, Medium E/I, High E/I) computed from the Ensembl GTF. The y-axis in all metagene plots represents the mean log2(Condition/Control) signal with ±1.96 SE confidence bands.

### Dual Partial Wave Spectroscopic (PWS) Imaging and Analysis

We performed Dual Partial Wave Spectroscopic (PWS) microscopy using a commercial inverted microscope (Leica DMIRB) equipped with a Hamamatsu Image-EM CCD camera (C9100-13) and a liquid crystal tunable filter (LCTF; CRi). The PWS microscope setup used in this study is described in detail by Almassalha et al^33^. All cells were maintained and imaged under physiological conditions (5% CO_2_ and 37°C) throughout imaging. Monochromatic spectrally resolved images were acquired from 500 to 700 nm at 1 nm intervals, with illumination provided by an Xcite-120 LED Lamp (Excelitas). The images were normalized by the incident light reflectance from the glass-media interface, using an independent reference image from a cell-free field of view to scale the reflectance.

In order to calculate chromatin packing scaling *D* for individual cells, we averaged Σ^2^ over all pixels across cell nuclei, then converted ⟨Σ^2^⟩ to *D* using custom MATLAB scripts^37^. In brief, the average nuclear spectral variance ⟨Σ^2^⟩ is multiplied by a correction factor of 2.43 to account for differences in internal reflection between microscopes. The resulting ⟨Σ^2^⟩ is then converted to *D* using our previously published computational model using input values. Typically, 100 to 200 cells were analyzed per condition to ensure robust statistical comparisons. Each treated cell population measured by PWS microscopy was compared to an untreated control population of the same cell type, which was plated on the same day with identical seeding density.

Briefly, PWS measures the spectral interference signal resulting from internal light scattering originating from nuclear chromatin. This is related to variations in the refractive index distribution (Σ) (extracted by calculating the standard deviation of the spectral interference at each pixel), characterized by chromatin packing scaling *D_n_*. *D_n_* was calculated using maps of Σ, as previously described^37^. Measurements were normalized by the reflectance of the glass medium interference, which allows us to obtain the interference signal directly related to the refractive index (RI) fluctuations within the cell. Although PWS is a diffraction-limited imaging modality, it can measure chromatin density variations because the RI is proportional to the local density of macromolecules, including DNA, RNA, and proteins. Therefore, Σ is proportional to chromatin density variations and can be characterized by the packing scaling behavior of chromatin with length scale sensitivity around 20-200 nm.

Pseudo-colored live cell PWS images were generated using Python by converting the Σ^2^ for each image pixel-wise to *D*_pixel,_ then mapping *D*_pixel_ values to a red colormap. *D*_pixel_ values ranging from 2 to 3 were visually represented, with higher values indicated by brighter red. This provides a spatial map of chromatin packing domain variations across the nucleus.

### Dynamic PWS Measurements

Dynamic PWS measurements were obtained as previously described^36^. Briefly, dynamic measurements (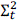, fractional moving mass (*m*_#_), and diffusion) are collected by acquiring multiple backscattered wide-field images at a single wavelength (550 nm) over time (acquisition time), to produce a three-dimensional image cube, where 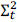 is temporal interference and t is time. Diffusion is extracted by calculating the decay rate of the autocorrelation of the temporal interference as previously described^36^. The fractional moving mass is calculated by normalizing the variance 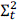 at each pixel. Using the equations and parameters supplied and explained in detail in Gladstein et al.^36^, the fractional moving mass is obtained by using the following equation to normalize 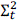 by *ρ*_$_, the density of a typical macromolecular cluster:

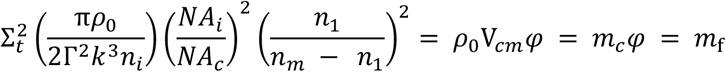

This normalization 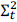 is equivalent to *m*_f_, which measures the mass moving within the sample. This value is calculated from the product of the mass of the typical moving cluster (*m_c_*) and the volume fraction of mobile mass (*φ*). *m*_’_ is obtained by *m_c_* = V_cm_*ρ*_0_, where V_cm_ is the volume of the typical moving macromolecular cluster. To calculate this normalization, we approximate *n*_)_ = 1.43 as the refractive index (RI) of a nucleosome, *n_m_* = 1.37 as the RI of a nucleus, *n*_&_ = 1.518 as the refractive index of the immersion oil, and *ρ*_0_ = 0.55 g *cm*^-3^ as the dry density of a nucleosome. Additionally, *k* = 1.57E5 cm^-1^ is the scalar wavenumber of the illumination light, and Γ is a Fresnel intensity coefficient for normal incidence. *NA_c_* = 1.49 is the numerical aperture (NA) of collection and *NA_i_* = 0.52 is the NA of illumination. As stated previously^36^, 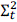 is sensitive to instrument parameters such as the depth of field, substrate refractive index, etc. These dependencies are removed through normalization with the proper pre-factor calculated above for obtaining biological measurements. It should also be noted that backscattered intensity is prone to errors along the transverse direction^36^. Due to these variations, these parameters are more accurate when calculating the expected value over each pixel.

### Single Molecule Localization Microscopy (SMLM) Staining, Imaging, and Analysis

#### EdU and H3K9me3 Dual-Label Staining and Imaging

Components of the Click-iT Plus EdU Cell Proliferation Kit for Imaging, Alexa Fluor 647 dye (Thermo Fisher Scientific, C10640), were stored according to the manufacturer’s instructions after proper reconstitution. Cells were seeded at 10,000 per well on No. 1 borosilicate-bottom, eight-well Lab-Tek chambered coverglass (Thermo Scientific, #155411) and allowed to adhere for 24 hours. EdU was added and incubated overnight to label cells throughout one complete cell cycle. Cells were then treated with auxin for 5 hours before being fixed for 10 minutes at room temperature in 4% paraformaldehyde (Electron Microscopy Sciences, #15710) in PBS (Gibco, #10010031), followed by three 5-minute washes in PBS. Quenching was done with freshly prepared 0.1% sodium borohydride (Sigma, #213462) in PBS for 7 minutes. Two more PBS wash steps were performed after quenching.

Permeabilization was done with blocking buffer composed of 3% bovine serum albumin (BSA) (Thermo Fisher Scientific, #BP671-10), 0.5% Triton X-100 (Thermo Fisher Scientific, #A16046-AE) in PBS for 1 hour. Samples were then incubated overnight at 4°C with anti-H3K9me3 antibody (Thermo Fisher Scientific, MA5-2567) in blocking buffer. Samples were then washed three times with a washing buffer composed of 0.2% BSA and 0.1% Triton X-100 in PBS.

Afterwards, samples were incubated with the corresponding goat antibody–dye conjugates, anti-mouse AF488 (ThermoFisher, A-11001) for 1 hour at room temperature with shaking. After incubation, samples were washed twice in PBS for 5 minutes, followed by imaging. Incorporation of EdU was visualized via click-chemistry-based secondary staining with AF647 fluorophore, which was performed exactly as outlined by the manufacturer.

#### EU Staining and Imaging

Components of the Click-iT RNA Alexa Fluor 594 Imaging Kit (Thermo Fisher Scientific, C10330) were stored according to the manufacturer’s instructions after proper reconstitution. Cells were seeded at 10,000 per well on No. 1 borosilicate-bottom, eight-well Lab-Tek chambered coverglass (Thermo Scientific, #155411) and allowed to adhere for 24 hours. Cells were then treated with auxin for 5 hours before being fixed for 10 minutes at room temperature in 4% paraformaldehyde (Electron Microscopy Sciences, #15710) in PBS (Gibco, #10010031), followed by three 5-minute washes in PBS. Incorporation of EU was visualized via click-chemistry-based secondary staining with AF5944 fluorophore, which was performed exactly as outlined by the manufacturer.

#### Single Molecule Localization Microscopy (SMLM) System Configuration

The STORM optical instrument was built on a commercial inverted microscope base (Nikon, Eclipse Ti-E with the perfect focus system). Two continuous lasers were used for illumination, one of which has five different wavelengths (405, 488, 532, 552, and 637 nm, Coherent, OBIS Laser Box) while the other is a green laser with 532 nm emission (Changchun New Industries Optoelectronics Tech. Co., Ltd., MGL-FN-532). The lasers are collimated with an average power at the sample of 3–10 kW/cm^3^. Images were collected via a ×100 1.49 numerical aperture (NA) objective (Nikon, SR APO TIRF) and captured on an electron-multiplying CCD camera (Andor, iXon Ultra 888). At least 10,000 frames with a 30 ms acquisition time were collected from each sample for each wavelength channel. All images were acquired using a total internal reflectance fluorescence illumination (TIRF) schema, which allowed for evanescent wave illumination of ∼200 nm slabs from the glass cell interface. Our focal plane was set within this depth to capture properly excited fluorophores that were in focus^2^.

#### Single Molecule Localization Microscopy (SMLM) Analysis

The raw image sequence was first pre-processed in ImageJ by subtracting its minimum-intensity projection and then applying a 5-pixel rolling-ball background subtraction to every frame. The preprocessed stack was imported into the ThunderSTORM plugin, where the camera was configured to match the acquisition settings: the pixel size of 130 nm (sensor pixel dimension divided by objective magnification), 18.6 photoelectrons per A/D count, a zero-count base level, and an EM gain of 240. For each cell, the nuclear ROI was drawn by Nuclei Outline in the GDSC plugin in ImageJ.

Final SMLM images were generated via standard reconstruction methods using the ThunderSTORM Fiji plugin with a maximum likelihood estimation of emission events with a 2D Gaussian PSF. Intensity was matched, so the ranges in all images were set to the mean of the control group. This was done to account for differences in localization events that may arise from the treatment effect^57^. Domains were identified using a previously developed pipeline that clusters localization events with Density-Based

Spatial Clustering of Applications with noise (DBSCAN) on reconstructed images. Parameters were optimized for the detection of chromatin packing domains^2^.

The association of H3K9me3 localizations with EdU-defined clusters was performed using the previously published multi-label SMLM pipeline^2^. EdU SMLM data served as the reference dataset for cluster identification. Clusters were defined by generating seed structures represented as convex polygon objects. For each cluster, the estimated radius was calculated based on the minimum enclosing circle corresponding to the convex polygon (i.e., the radius of the smallest circumscribed circle). To enable efficient spatial querying, a cKDTree was constructed from the H3K9me3 estimated localization coordinates. Each EdU cluster centroid was then used as a query point against the H3K9me3 cKDTree. A radial search was performed using a threshold distance of 1.5 × the estimated cluster radius to identify H3K9me3 localizations within the neighborhood of the reference cluster. All H3K9me3 points falling within this search radius were assigned the corresponding cluster ID in the output data frame. In cases where a localization fell within overlapping search regions of multiple clusters, assignment was resolved based on the shortest Euclidean distance to the respective cluster centroids. The resulting dataset was subsequently partitioned by cluster size category, and visualization was performed using the Python library matplotlib.

#### Phenomenological Model of Domain Formation

To study how crosslinking mechanisms play a role in packing domain stability and function, we extended and used a previously developed phenomenological model of domain formation that captures the physical interplay between chromatin crowding, transcriptional activity, and polymer folding dynamics. This model is grounded in three core principles. Namely, (1) loop returns (originating from transcription or cohesion-mediated loop extrusion) generate localized chromatin condensates, (2) macromolecular crowding influences the ability of enzymes and regulatory proteins to access and modify chromatin, and (3) transcriptional efficiency, which exhibits a non-monotonic dependence on local chromatin density, arises from the trade-off between diffusivity and reaction free energy.

Within this model, chromatin is treated as a flexible polymer confined within a nuclear subvolume that is heterogeneously packed within the nucleus and forms packing domains. Packing domains are assumed to follow a fractal polymer scaling law, where the local chromatin volume fraction (Eq.1) decays radially as a function of the center of the packing domain

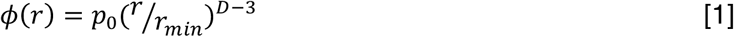

Where *r_min_*_*_ = 10 *nm* is the minimal chromatin chain element length, *p*_0_ is the chromatin volume concentration at the core of a packing domain (*r* = 0), and *D* is the domain’s packing scaling exponent, which governs how chromatin mass scales with domain size. This formulation enables quantification of both crowding and spatial organization within packing domains.

The model consists of a system of coupled ordinary differential equations (ODEs) that describe the temporal evolution of chromatin states.

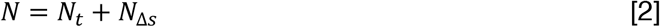

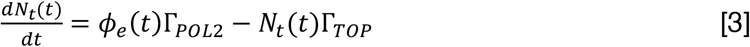

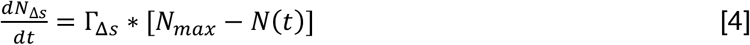

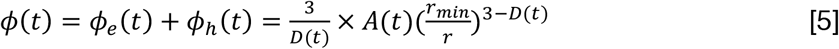

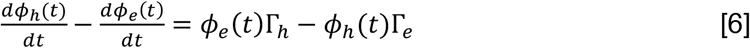

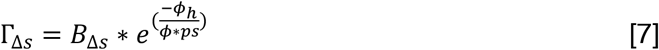

Here, *N_t_*(*t*) is the fraction of chromatin engaged in transcriptional loop returns, *N*_Δ*s*_(*t*) corresponds to the fraction of chromatin undergoing entropic (non-enzymatic) returns, *φ_h_*(*t*) and *φ_e_*(*t*) describe the chromatin volume fractions of heterochromatin and euchromatin, and *D*(*t*) is the time-varying packing scaling exponent describing how the polymer scales with size. Returns from either transcriptional mechanisms or entropic crosslinking, each associated with distinct biophysical feedback. Eq. 3 governs the time evolution of transcriptional returns, controlled by two rate constants: Γ*_POL2_*, representing RNA Polymerase II activity, and Γ*_TOP_*, representing topoisomerase-mediated loop relaxation. These rate constants exhibit non-monotonic dependencies on local chromatin composition, particularly the balance between euchromatin and heterochromatin volume fractions. Enzymatic activity is modeled as a function of local crowding, where the return formation reflects the trade-off between chromatin accessibility and chromatin density. Accessibility is further modulated by chromatin folding via a mechanistic function *f*(*D*(*t*), *p_c_*, *p*_0_, *r_min_*, *r*) where *p_c_* defines the enzyme-specific crowding tolerance. A complete breakdown of the model and equations, including explicit equations for the rate constant and folding function, can be found in Almassalha et al^4^.

## Statistical Analysis

Significance was calculated by an unpaired *t*-test with Welch’s correction applied in GraphPad Prism 10.1.1 and Python. For all figures, a *p*-value less than 0.05 was considered significant. Statistical significance levels are denoted as follows: ns; \**p* < 0.05; \*\**p* < 0.01; *** *p* < 0.001; **** *p* < 0.0001. For RNA sequencing data, differential gene expression was evaluated using DESeq2, and significance was determined using false discovery rate (FDR)-adjusted p-values. Genes with adjusted p < 0.05 were considered significantly differentially expressed. All experiments were independently repeated at least twice with biological duplicates to ensure reproducibility of results.

## Declarations

### Ethics Approval and Consent to Participate

Not applicable.

### Consent for Publication

Not applicable

### Availability of Data and Materials

The custom scripts for data analysis and visualization are available in the Backman Lab GitHub repository (https://github.com/BackmanLab/HP1a-Paper). The RNA sequencing data that support the findings of this study have been submitted to the NCBI Sequence Read Archive (SRA) under BioProject accession number PRJNA1516088. All other data are included in the main manuscript and/or supplementary information.

## Authors’ Contributions

T.K. wrote the main manuscript text; T.K. and B.J. conducted RNA-seq experiments; T.K., P.C.G, and L.M.C. conducted RNA-seq analysis; T.K. conducted PWS imaging and analysis; N.A. and C.V.A. conducted SMLM imaging and analysis; P.C.G. conducted modeling experiments and analysis; E.P.L. conducted flow cytometry experiments; T.K. conducted confocal microscopy imaging; T.K., P.C.G., N.A., C.L.D., L.M.A., I.S., and V.B. contributed to project conceptualization, troubleshooting, and manuscript development through intellectual discussions and feedback.

## Acknowledgements

We would like to thank Dr. Andrew D. Stephens for sharing the U2OS HP1α-AID cell line with us. We would also like to thank Dr. Jane Frederick for their helpful discussions during the experimental design and initial manuscript preparation. We thank the Robert H. Lurie Comprehensive Cancer Center of Northwestern University in Chicago, IL, for the use of the Flow Cytometry Core Facility, which provided flow cytometric analysis training and use of the BD LSRFortessa Cell Analyzer FACSymphony S6 SORP system. We thank the Biological Imaging Facility at Northwestern University (RRID: SCR_017767) for providing training and use of the Nikon SoRa Spinning Disc confocal microscope. This work was supported by the Northwestern University NUSeq Core Facility, where bulk RNA-Seq library preparation was performed.

This research was supported in part through the computational resources and staff contributions provided by the Genomics Compute Cluster, which is jointly supported by the Feinberg School of Medicine, the Center for Genetic Medicine, and Feinberg’s Department of Biochemistry and Molecular Genetics, the Office of the Provost, the Office for Research, and Northwestern Information Technology. The Genomics Compute Cluster is part of Quest, Northwestern University’s high-performance computing facility, with the purpose of advancing genomics research.

ChatGPT-5 was utilized to assist in writing and debugging some data visualization scripts in Python and R. The authors thoroughly reviewed and verified the accuracy of all code and content, maintaining full responsibility for the manuscript’s integrity.

