## Supplementary Information for "Heterochromatin protein 1α acts as an essential stabilizer of chromatin packing domains"

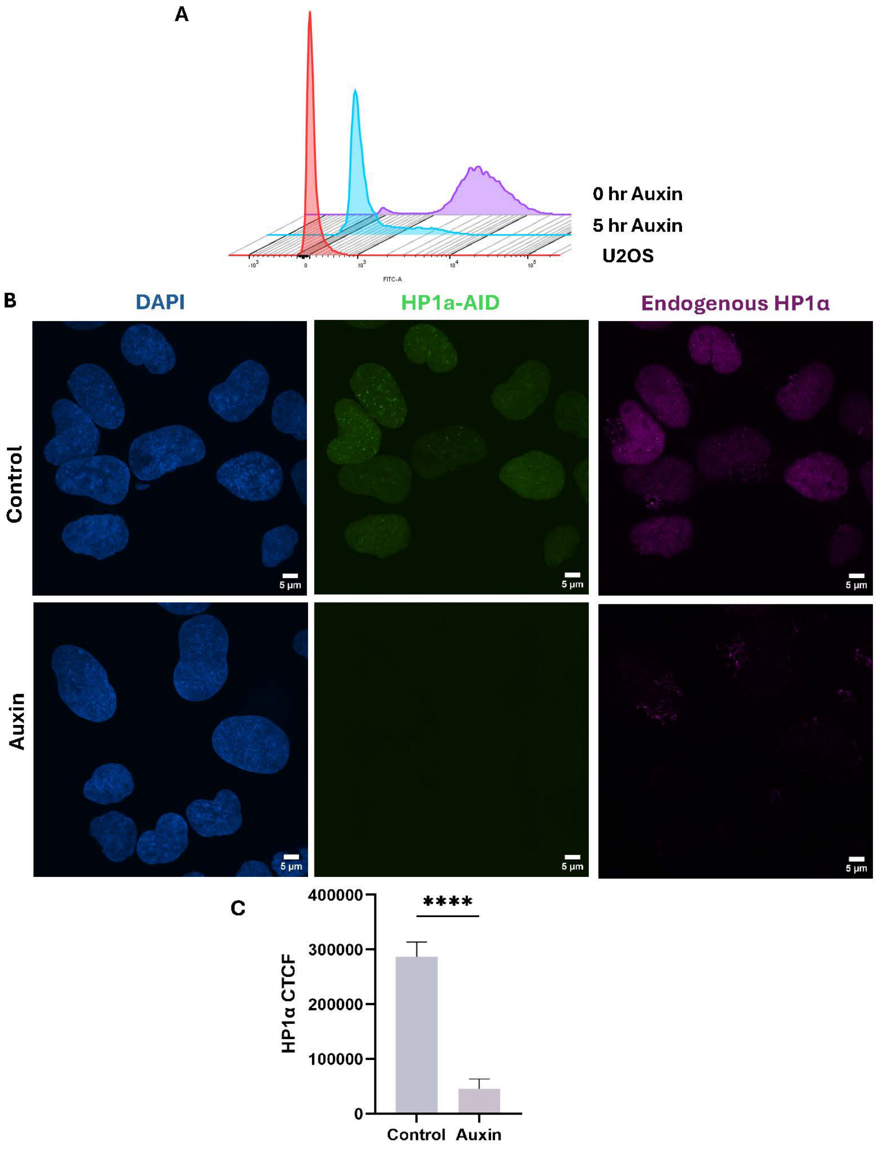


**Supplementary Figure 1: Auxin treatment sufficiently degrades HP1α within 5 hours**

A. Flow cytometric analysis shows that auxin treatment sufficiently degrades HP1α in U2OS-HP1α-AID cells within 5 hours.

B. Immunostaining of HP1α tagged with AID and endogenous HP1α in U2OS-HP1α-AID cells. Blue labels nuclei with DAPI staining, green labels HP1α-AID, and magenta labels endogenous HP1α. Scale bars = 5 µm.

C. Bar graphs for Corrected Total Cell Fluorescence (CTCF) of endogenous HP1α in control and auxin-treated U2OS-HP1α-AID cells in confocal imaging. Significance was calculated by an unpaired t-test with Welch's correction applied, with ****$P<0.0001$.

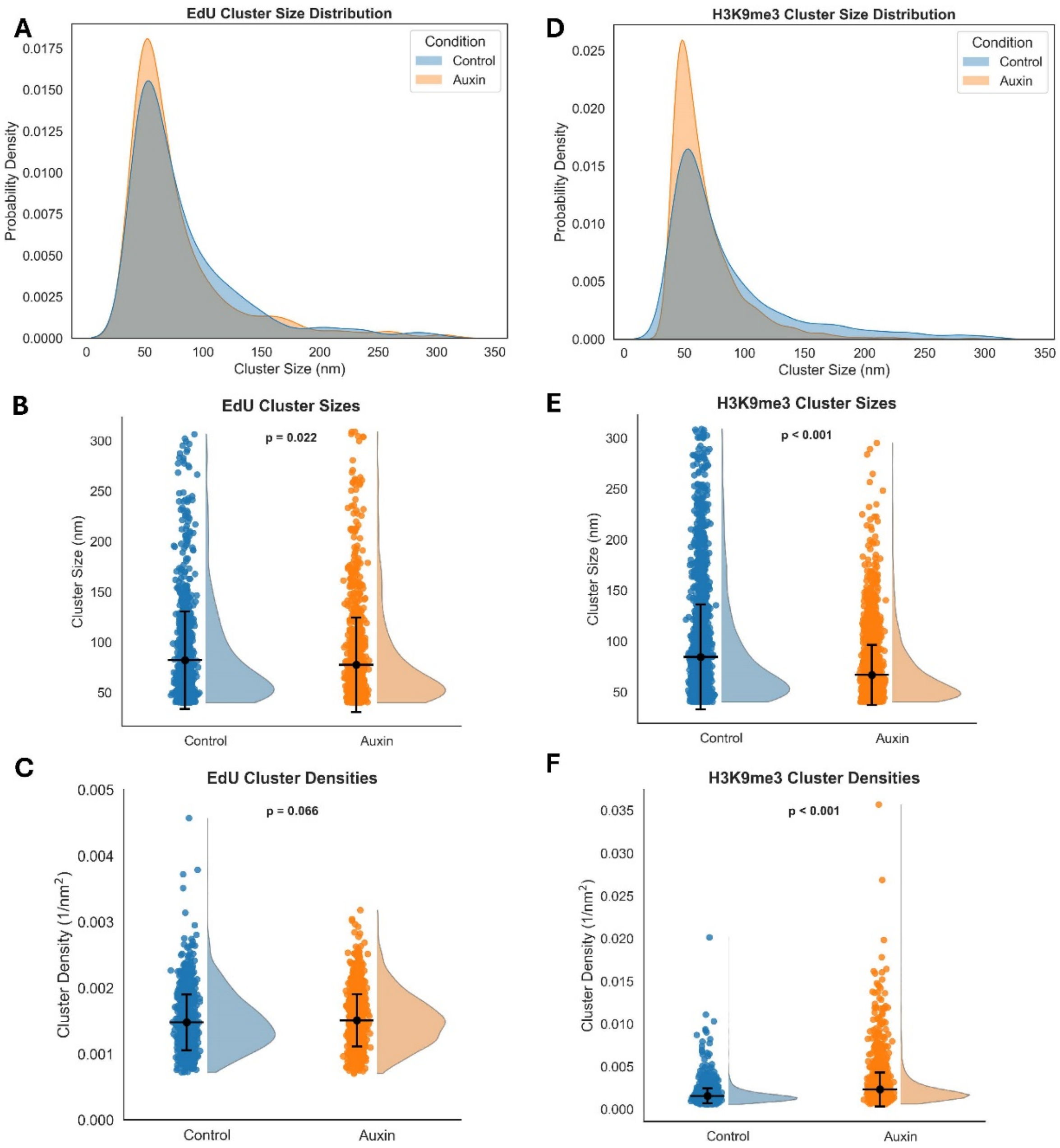


**Supplementary Figure 2: Additional quantifications and analyses of single molecule localization microscopy data**

A. Probability density function of EdU cluster sizes in control and auxin-treated U2OS-HP1α-AID cells, with auxin-treated cells having a slight increase in the number of smaller EdU-defined clusters relative to the control.

B. Quantification of EdU cluster sizes in control and auxin-treated U2OS-HP1α-AID cells.

C. Quantification of EdU cluster densities in control and auxin-treated U2OS-HP1α-AID cells.

D. Probability density function of H3K9me3 cluster sizes in control and auxin-treated U2OS-HP1α-AID cells, with auxin-treated cells having an increased number of smaller H3K9me3-defined clusters relative to the control.

E. Quantification of H3K9me3 cluster sizes in control and auxin-treated U2OS-HP1α-AID cells.

F. Quantification of H3K9me3 cluster densities in control and auxin-treated U2OS-HP1α-AID cells.

For B-C, E-F: Significance was calculated by an unpaired t-test with Welch's correction applied. Calculated p-values are denoted on the graphs unless p-value < 0.001.

**A**

| Parameter | Mature Domain | Nascent Domain |
|---|---|---|
| Starting Packing Scaling $D_{initial}$ | 2.69 | 2.2 |
| Optimal Loop Capacity within the Volume $N_{max}$ | 0.0289 | 0.0131 |
| Starting Fraction of Chromatin in Returns Due to Transcription $N_t(0)$ | $N_{max} * 0.7$ | $N_{max} * 0.1$ |
| Starting Fraction of Chromatin in Returns Due to Entropic Stabilization/Crosslinking $N_s(0)$ | $N_{max} * 0.3$ | $N_{max} * 0.9$ |
| Starting Packing Efficiency $P_0(0)$ | 0.73 | 0.44 |
| Starting Heterochromatic Chromatin Volume Concentration $\phi_h(0)$ | 0.2 | 0.005 |

**Supplementary Figure 3: Parameters that define mature and nascent packing domains in phenomenological model of domain formation**

A. List of parameters that classify mature and nascent packing domains, including starting packing scaling, optimal loop capacity within the volume, starting fraction of chromatin in returns due to transcription, starting fraction of chromatin in returns due to entropic stabilization/crosslinking, starting packing efficiency, and starting heterochromatic chromatin volume concentration.

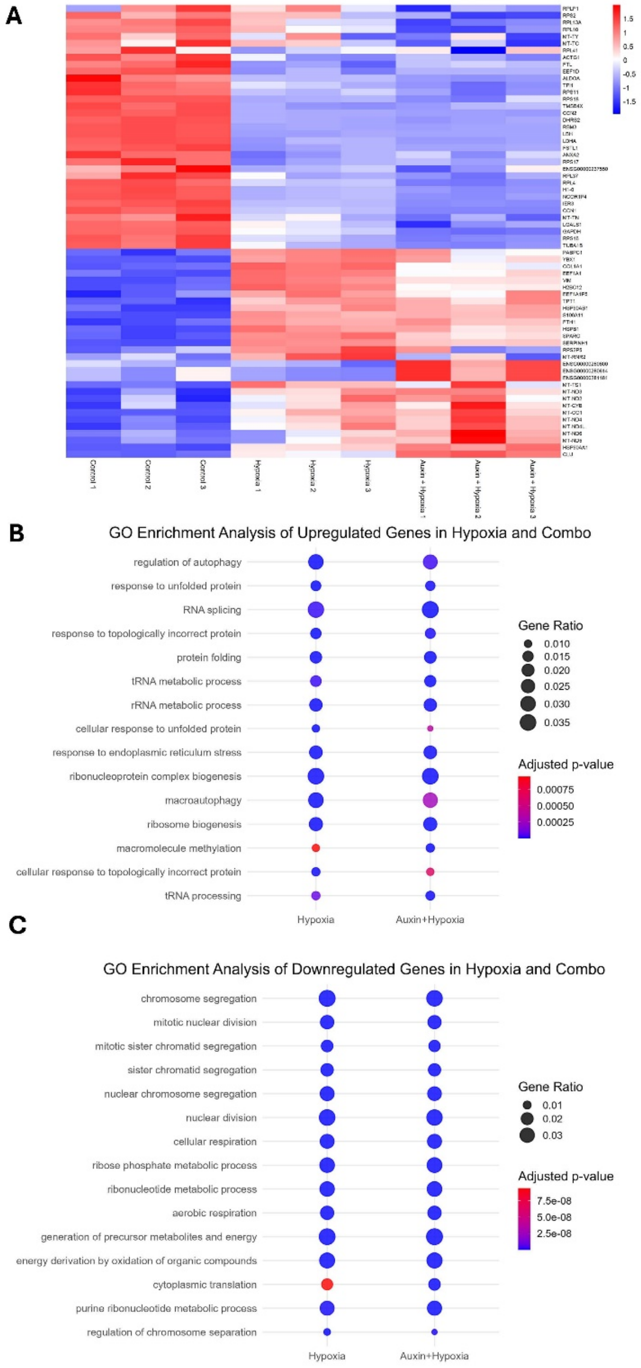


**Supplementary Figure 4: HP1α-depleted cells that were exposed to hypoxia had similar transcriptional profiles compared to hypoxia**

A. Heatmap of the top 65 differentially expressed genes in all three replicates of Control, Hypoxia, and Auxin + Hypoxia conditions

B. Top 15 upregulated Gene Ontology (GO) biological processes (BP) terms in hypoxia and auxin + hypoxia treatments.

C. Top 15 downregulated Gene Ontology (GO) biological processes (BP) terms in hypoxia and auxin + hypoxia treatments.

For B-C, the sizes of the circles represent the proportion of differentially expressed genes that fall into a specific biological pathway, while the colors represent the statistical significance (adjusted p-value).

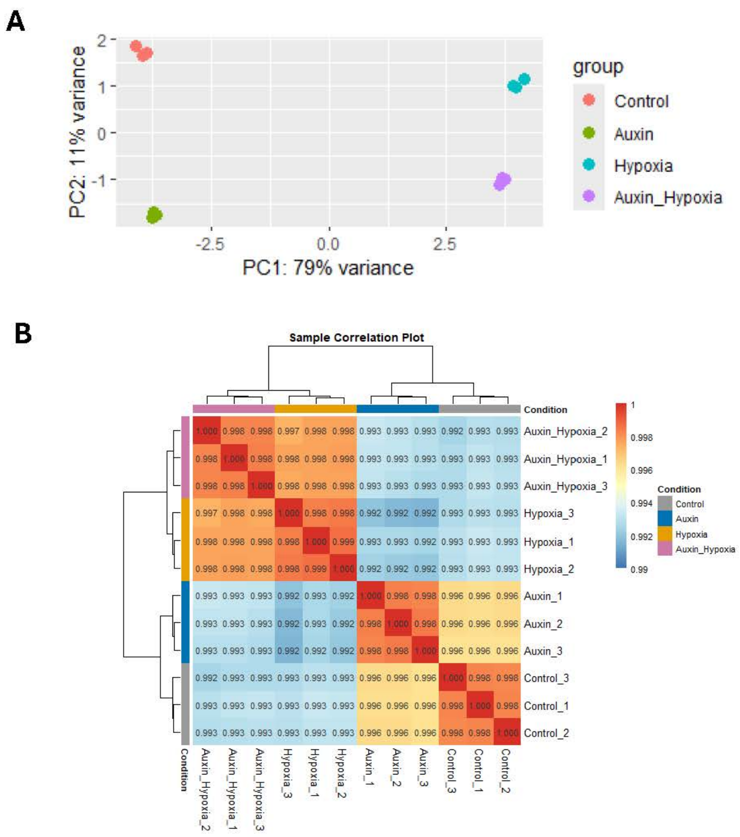


**Supplementary Figure 5: Additional analyses to ensure quality of bulk RNA sequencing data**

A. Principal component analysis (PCA) of normalized RNA sequencing data. Samples distinctly cluster based on their experimental group and demonstrate clear separation along the first two principal components, indicating distinct transcriptional profiles among groups.

B. Pearson correlation heatmap of normalized RNA sequencing data. All three biological replicates of each treatment exhibited strong positive correlations and clustered according to experimental condition. Higher within-group relations indicate high reproducibility and distinct gene expression patterns among groups.

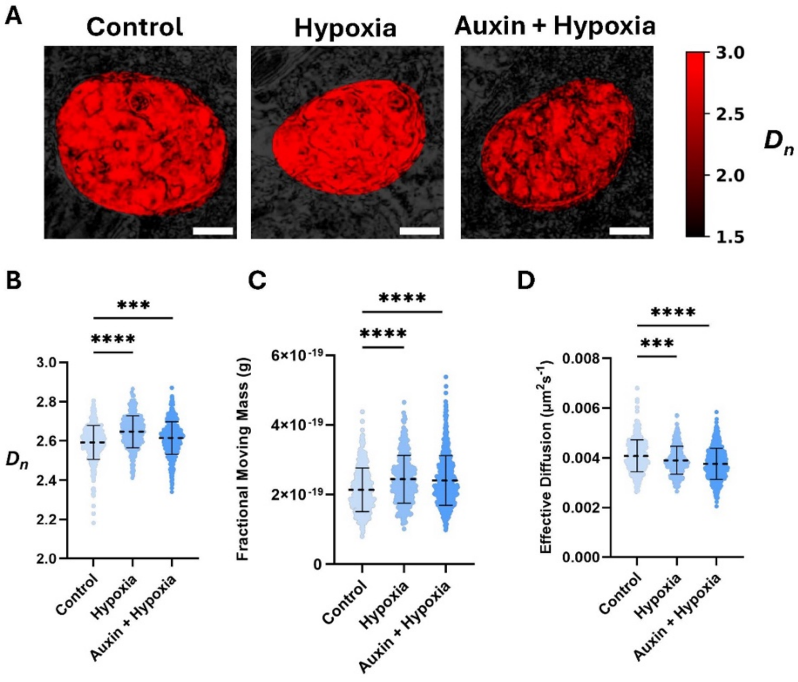


**Supplementary Figure 6: HP1α-depleted cells can generate nascent packing domains in response to a stressor**

A. Representative images of U2OS-HP1α-AID cells that were untreated, exposed to 24 hours of hypoxia, and 5 hours of auxin treatment before being exposed to 24 hours of hypoxia. Scale bars = 5 μm.

B. Violin plots for chromatin packing scaling ($D_n$) in U2OS-HP1α-AID cells that were untreated, exposed to 24 hours of hypoxia, and treated with 5 hours of auxin before being exposed to 24 hours of hypoxia.

C. Violin plots for fractional moving mass in U2OS-HP1α-AID cells that were untreated, exposed to 24 hours of hypoxia, and treated with 5 hours of auxin before being exposed to 24 hours of hypoxia.

D. Violin plots for diffusion coefficient in U2OS-HP1α-AID cells that were untreated, exposed to 24 hours of hypoxia, and treated with 5 hours of auxin before being exposed to 24 hours of hypoxia.

For B-D: Data were compiled from three independent biological replicates. Significance was calculated by an unpaired *t*-test with Welch's correction applied, with ***P<0.001 and ****P<0.0001. (Control n=351; Hypoxia n=324; Auxin and Hypoxia n=558)

**A**

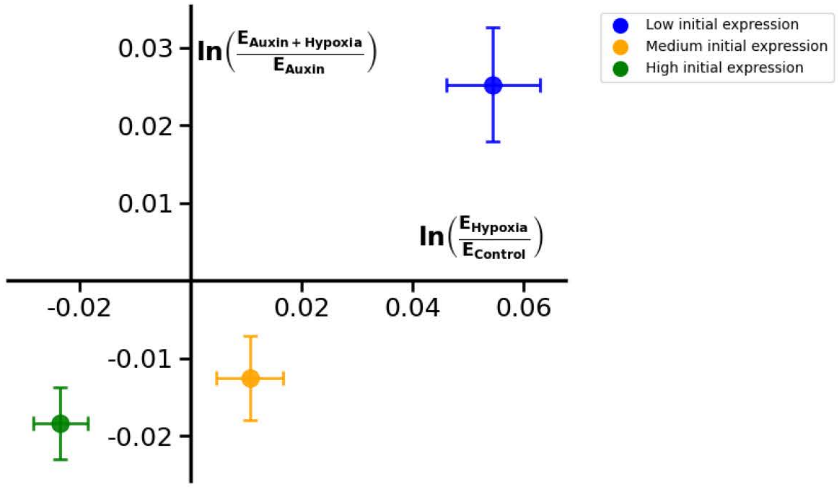


**B**

| | Group | n_genes | slope | intercept | R2 | p_value | std_y |
|---|---|---|---|---|---|---|---|
| **0** | Medium initial expression | 2168 | 0.7726 | -0.0208 | 0.7382 | 0.0 | 0.2548 |
| **1** | High initial expression | 2168 | 0.8197 | 0.0008 | 0.7530 | 0.0 | 0.2154 |
| **2** | Low initial expression | 2169 | 0.7429 | -0.0153 | 0.7237 | 0.0 | 0.3426 |

**Supplementary Figure 7: Additional analyses of bulk RNA sequencing data**

A. Hypoxia-responsive genes were identified based on their significance relative to the control. These genes were then binned based on their initial expressions, and the average response for each group was plotted to determine whether the presence or absence of HP1α alters the hypoxia-responsive genes' abilities to respond to the external stressor.

B. The table quantifies the number of genes in each binned group of significantly expressed hypoxia-responsive genes, slopes, intercepts, $R^2$ values, p-values, and standard deviations on the y-axis.
